# HDAC Inhibition Sensitizes Pancreatic Tumors to DNA Damage by Global Redistribution of the Transcriptional Machinery

**DOI:** 10.64898/2026.05.18.726071

**Authors:** Gaoyang Liang, Hung V.-T. Nguyen, Jonathan Zhu, Hervé Tiriac, Hadiqa Zafar, Daniel Y. Cao, Gabriela Estepa, Dylan C. Nelson, Yang Dai, Tae Gyu Oh, Christopher Liddle, Ruth T. Yu, Tony Hunter, Dannielle Engle, Reuben Shaw, Andrew M. Lowy, Weiwei Fan, Morgan L. Truitt, Annette R. Atkins, Jeremiah A. Johnson, Michael Downes, Ronald M. Evans

## Abstract

The DNA damage response (DDR) is critical for pancreatic ductal adenocarcinoma (PDAC) development and therapeutic responses, including to genotoxic agents. While epigenetic modulators have been shown to contribute to the DDR, how chromatin regulation dictates responses to DNA damage in PDAC remains incompletely understood. Here, we identify Class I histone deacetylases (HDACs) as critical regulators of the DDR. HDAC1/2 directs the genomic distribution of H3K27ac, ensuring sufficient BRD4 and RNA polymerase II (Pol II) occupancy at DDR gene promoters. HDAC inhibition by entinostat shifts the balance of H3K27 acetylation preferentially towards intergenic regions, diverting BRD4 and Pol II from promoters, thereby suppressing DDR gene expression. In line with this, HDAC inhibition heightens DNA damage and sensitizes PDAC to diverse DNA-damaging and DDR-targeting agents. Since the clinical development of HDAC inhibitors has been limited by systemic toxicity, we developed bottlebrush prodrug (BPD) nanoparticles for tumor-selective entinostat delivery. Entinostat-BPD achieved tumor-specific HDAC inhibition while displaying potent efficacy and reduced systemic toxicity. These findings reveal an HDAC-dependent DDR vulnerability and offer combinational and precision targeting strategies to facilitate clinical translation and improve PDAC patient outcomes.

**SIGNIFICANCE STATEMENT:** The ability of tumor cells to tolerate DNA damage limits the efficacy of many anticancer therapies. Our study reveals that pancreatic cancer cells enforce this resistance by sustaining expression of DNA damage response (DDR) genes through Class I histone deacetylases (HDACs). HDACs maintain genome-wide acetylation patterns required for efficient recruitment of the transcriptional machinery to DDR genes. Pharmacological HDAC inhibition disrupts this process and sensitizes pancreatic cancer cells to diverse DNA-damaging agents. To overcome systemic toxicity that limits translational potential, we further establish a bottlebrush prodrug nanoparticle platform that enables tumor-selective HDAC inhibition. Given the central role of the DDR in cancer, targeting HDAC-mediated DDR regulation through drug combinations and precision delivery may have broad therapeutic relevance across cancer types.

## INTRODUCTION

Pancreatic ductal adenocarcinoma (PDAC) is among the deadliest cancers, with limited therapeutic options and poor survival (1, 2). Like many cancers, current therapies for PDAC rely heavily on DNA-damaging agents, including platinum-based crosslinkers (oxaliplatin, cisplatin), topoisomerase inhibitors (irinotecan), and antimetabolites (gemcitabine, 5-florouracil or 5-FU) (3–6). These agents generate DNA lesions that, if unrepaired, accumulate to induce cytotoxicity and senescence (7). Yet their efficacy in PDAC remains modest, reflecting profound resistance driven by both tumor-intrinsic and -extrinsic mechanisms (8–10). Notably, patients with mutations in DNA damage response (DDR) components, such as BRCA1, BRCA2, or PALB2, exhibit improved responses to genotoxic agents and DDR inhibitors (11–13), underscoring the dependency of PDAC on intact DDR pathways to withstand genotoxic stress. Elucidating the mechanisms that sustain this dependency may reveal new therapeutic opportunities (14–17).

Chromatin regulation has been shown to contribute to the DDR by coordinating the recruitment of DDR factors, influencing repair pathway choice, facilitating restoration, and modulating gene expression (18–21). In particular, Class I histone deacetylases (HDACs) are recruited to double strand breaks (DSBs), where they facilitate non-homologous end joining (NHEJ) through deacetylation of histones (e.g., H3K56ac) and repair factors (e.g., Ku70) (22, 23). While HDACs have been classically viewed as transcriptional repressors that compact chromatin (24, 25), HDACs have also been implicated in supporting the DDR transcriptionally, as HDAC inhibition suppresses DDR gene expression and sensitizes multiple cancer types to genotoxic agents (21, 26–29). Although growing evidences link HDACs to active transcription (30–38), the molecular mechanisms by which HDACs support transcription of DDR genes remains poorly understood.

Despite the correlations of HDACs with poor prognosis and the anti-tumor activity of HDAC inhibitors (HDACis) across cancer models (26, 28, 29, 39), including PDAC (38, 40–43), the clinical utility of HDACis has been largely confined to a few hematologic malignancies, with limited efficacy in solid tumors (29, 44–46). This limitation arises from narrow or often absent therapeutic windows, as systemic inhibition of HDACs disrupts essential physiological functions alongside tumor suppression. One approach for overcoming this limitation is to identify combinational strategies that amplify the efficacy of HDACis at their tolerable doses (29, 38, 42, 43, 47). Tumor-selective delivery of HDACis is another promising approach. Bottlebrush prodrug (BPD) nanoparticles have emerged as an effective platform that enables tumor-enriched accumulation and tumor-selective drug release, thereby reducing systemic exposure and toxicity. While the BPD platform has been successfully used to achieve tumor-selective delivery of chemotherapeutics and BET inhibitors in other cancers (48, 49), its utility in targeting PDAC or delivering HDACis has not been explored.

Here, we reveal that Class I HDACs support DDR gene transcription in PDAC and establish combinational and BPD nanoparticle-based strategies to potentiate HDACi therapy. Through integrated transcriptomic and epigenomic analyses, we show that HDACs maintain a proper balance of genome-wide chromatin acetylation distribution, which is required for the localization of the transcriptional machinery, including BRD4 and Pol II, to facilitate DDR gene expression. HDAC inhibition disrupts this balance, suppresses DDR gene expression, and correspondingly enhances PDAC tumor vulnerability to DNA-damaging agents. Furthermore, we demonstrate that tumor-selective delivery of HDACi through BPD nanoparticles achieves anti-tumor efficacy, mitigates systemic toxicity, and potentiates DNA-damaging therapies, presenting a tractable strategy to improve PDAC treatment.

## RESULTS

### Class I HDACs Support the DDR Transcriptional Program in PDAC

Given the central role of epigenetic regulation in driving tumor dependency and therapeutic resistance (50, 51), we investigated the contributions of Class I HDACs to PDAC. Human and mouse PDAC cells were treated with the Class I HDAC inhibitor entinostat (5 µM, 24 h) and subjected to genome-wide transcriptomic analysis. Entinostat treatment upregulated and downregulated over 3,000 genes, respectively (Fig. 1*A* and *B*). Significantly suppressed biological processes included cell division, translation, DNA repair, and the DNA damage response (*SI Appendix*, Fig. S1*A*), while preferentially upregulated processes included apoptosis and autophagy (*SI Appendix*, Fig. S1*B*). Notably, downregulated DDR genes — particularly DNA repair genes — encompassed key regulators of multiple pathways sensing and repairing diverse forms of DNA damage (Fig. 1*A–D*). The transcriptional effects of entinostat treatment could largely be recapitulated by shRNA-mediated co-depletion of HDAC1 and HDAC2, while depletion of HDAC1 or HDAC2 alone produced substantially weaker effects (*SI Appendix*, Fig. S1*C*–*E*), consistent with their known functional redundancy (24, 52). On the other hand, depletion of HDAC3 had minimal impact on entinostat-sensitive programs (*SI Appendix*, Fig. S1*C*–*E*), likely reflecting its unique complex association and distinct function (24, 52). In line with the role of Class I HDACs in sustaining the DDR transcriptional program, entinostat treatment led to accumulation of the DNA damage marker γH2A.X, along with increased histone acetylation from HDAC inhibition (Fig. 1*E* and *F*), further supporting Class I HDACs as critical for the DDR in PDAC.

**Fig. 1.**
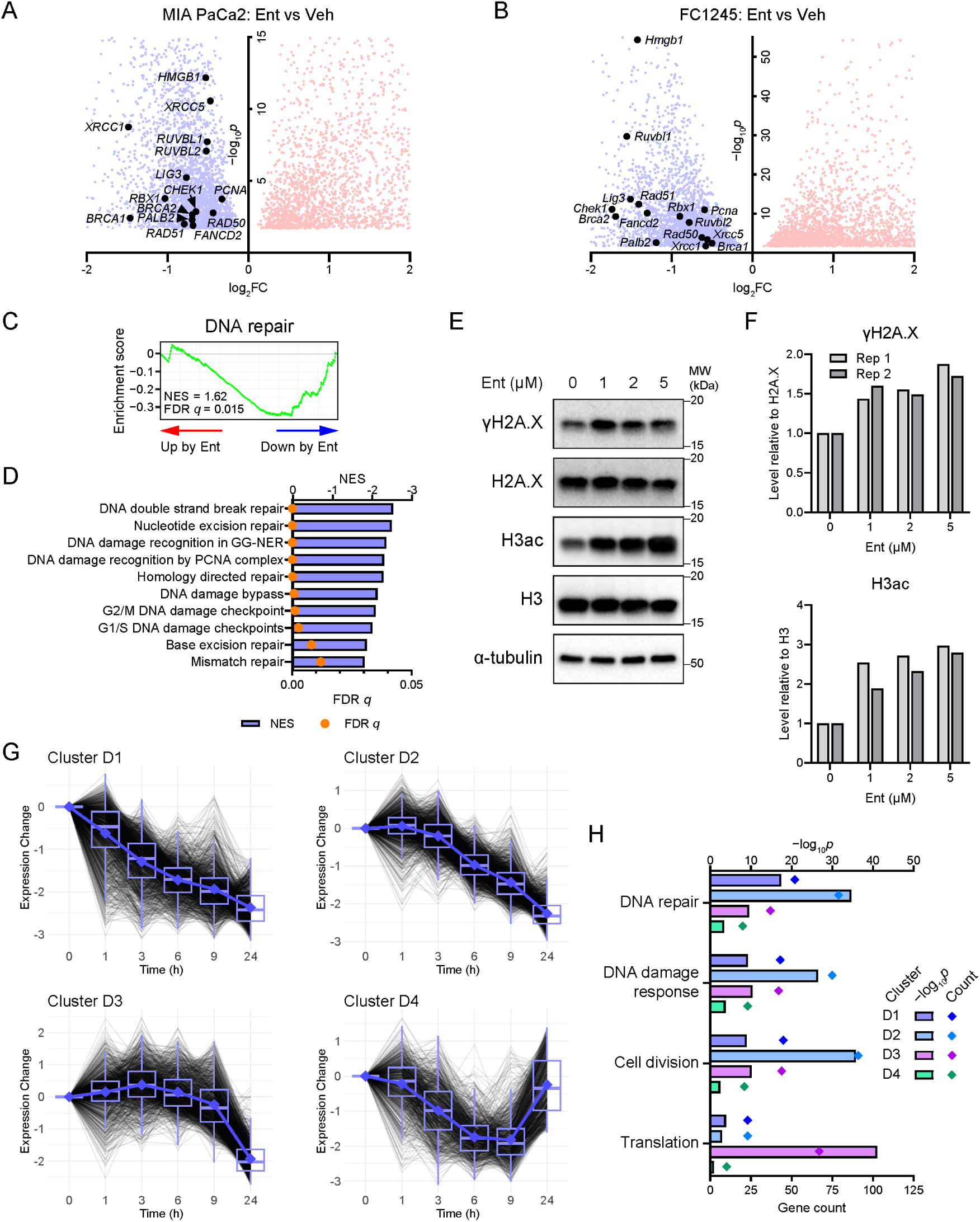
Entinostat treatment suppresses DDR pathways and induces DNA damage in PDAC. (*A* and *B*) Volcano plots showing RNA-seq results of differentially expressed genes between entinostat (Ent, 5 µM, 24 h) and vehicle (Veh) treatment in human MIA PaCa2 (*A*) and mouse FC1245 (*B*) PDAC cells. 3602 (MIA PaCa2) and 3543 (FC1245) genes are significantly upregulated, and 3478 (MIA PaCa2) and 3158 (FC1245) genes downregulated with representative DDR-related genes highlighted. FC, fold change. *n* = 2 independent cell samples. (*C*) GSEA enrichment plot showing DNA repair hallmark enriched in Ent-downregulated genes in FC1245. NES, normalized enrichment score; FDR, false discovery rate; Up, upregulation; Down, downregulation. (*D*) DDR-related gene ontology (GO) terms enriched in Ent-downregulated genes from GSEA. GG-NER, global genome nucleotide excision repair; PCNA, proliferating cell nuclear antigen. (*E* and *F*) Representative Western blots (*E*) and quantifications (*F*) showing increased γH2A.X and H3ac (normalized to histone H2A.X and H3, respectively) under graded Ent treatments. *n* = 2 independent experiments. (*G*) 4 gene clusters with downregulation trends (D1–D4) identified by temporal expression analysis in FC1245 cells with 0–24 h treatment of Ent (5 µM). *n* = 3 independent cell samples. (*H*) Enrichment of selected GO terms in downregulated clusters.

To assess whether the suppression of DDR genes represents an immediate, direct effect of HDAC inhibition and to understand how this relates to other HDAC-regulated biological processes, we profiled the transcriptome dynamics over 1–24 h of entinostat treatment. Temporal analysis identified 8 gene clusters with distinct expression patterns: 4 clusters primarily upregulated (U1–U4; *SI Appendix*, Fig. S2*A* and *B*) and 4 downregulated (D1–D4; Fig. 1*G* and *SI Appendix*, Fig. S2*A*). Among the downregulated clusters, immediate (D1, down starting at 1 h) and early (D2, down starting at 6 h) responses together accounted for ∼57% of affected genes, while late responses (D3, down at 24 h only) accounted for ∼25% (Fig. 1*G* and *SI Appendix*, Fig. S2*C, D*). In addition, there was a subset of downregulated genes (D4, ∼17%) that showed initial suppression (persisting up to 9 h) followed by compensation after prolonged treatment (back up at 24 h) (Fig. 1*G* and *SI Appendix*, Fig. S2*C* and *D*). Notably, genes involved in DNA repair and DDR, along with those regulating cell division, were highly enriched in D1 and D2 (Fig. 1*H*), suggesting a direct effect of HDAC inhibition. In contrast, translation-related genes were predominantly enriched in D3 and downregulated later (Fig. 1*H*), likely reflecting a secondary effect from earlier transcriptional changes. Among the biological processes enriched in the upregulated genes, apoptosis-related genes were enriched across similarly identified immediate, early, and late responses (U1–U3; *SI Appendix*, Fig. S2*B*, *E*–*G*), whereas autophagy-related genes were predominantly induced late (U3; *SI Appendix*, Fig. S2*B*, *E*–*G*) in line with findings that autophagy can be an adaptive pro-survival response (53, 54). Together, these transcriptomic analyses demonstrate that Class I HDACs sustain the DDR transcriptional program in PDAC, likely through direct regulation.

### HDACs Localize at DDR Gene Promoters

To elucidate how HDACs directly regulate transcription, we mapped the genome-wide occupancy of HDAC1 and HDAC2 in PDAC cells by the cleavage under targets and release using nuclease (CUT&RUN) assays (55). We identified a total of 19,005 genomic sites bound by HDAC1 and/or HDAC2, across promoters (∼32%), intergenic (∼30%), and genic (∼38%) regions (Fig. 2A and *B*). HDAC1 and HDAC2 displayed highly similar occupancy patterns at these sites (Fig. 2*B* and *SI Appendix*, Fig. S3), consistent with their shared biochemical and functional properties (52). Interestingly, these HDAC1/2-bound sites were co-occupied by active histone marks and transcription machinery, including H3K4me3 at promoters and H3K27ac, BRD4 (a key acetyl-binding transcriptional effector), and RPB1 (the major Pol II subunit) genome-wide (Fig. 2*B* and *SI Appendix*, Fig. S3), supporting a role of HDAC1/2 in active transcription.

**Fig. 2.**
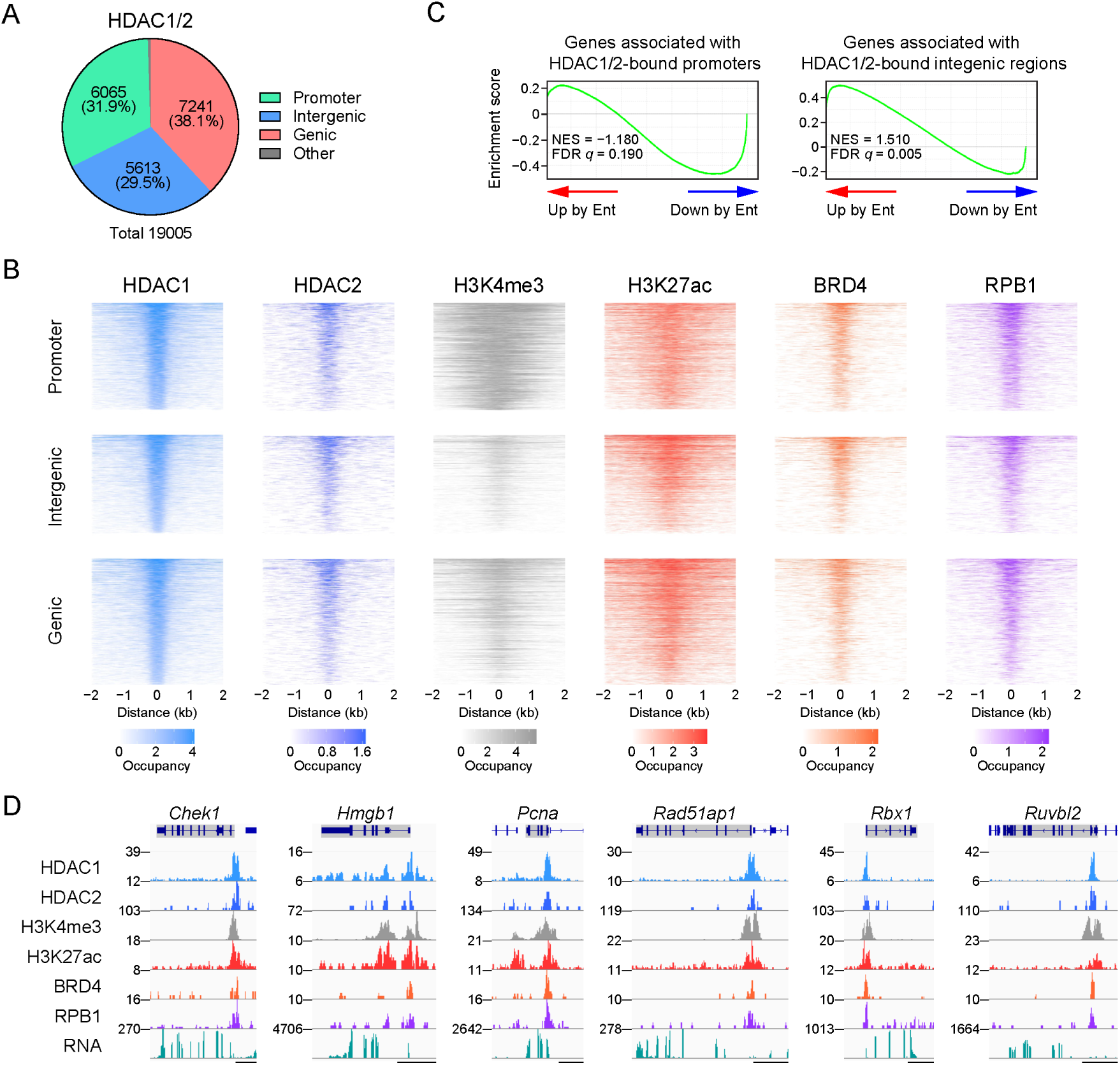
HDAC1/2 localize at DDR gene promoters with transcription machinery. (*A*) Numbers and percentages of annotated peaks bound by HDAC1 and/or HDAC2 in FC1245 as detected by CUT&RUN. (*B*) Heatmaps of HDAC1, HDAC2, H3K27ac, BRD4, and RPB1 occupancy at HDAC1/2-bound regions. Occupancy is log_2_-transformed. (*C*) GSEA plots showing correlations of Ent-induced upregulation (Up) and downregulation (Down) with gene sets linked to the top 1000 HDAC1/2-bound promoters or intergenic regions. (*D*) Genome browser tracks of representative DDR genes showing the localizations of HDAC1/2 with transcription machinery at promoters along with RNA-seq data. Scale bar, 5 kb.

To link HDAC1/2 occupancy with transcriptional outcomes, we assigned the identified HDAC1/2-bound sites to their nearest genes and examined their expression upon entinostat treatment. Genes with HDAC1/2-bound promoters were preferentially downregulated by entinostat (Fig. 2*C*, left) and enriched for DNA repair, DDR, and other HDACi-suppressed pathways (*SI Appendix*, Fig. S4*A*). Promoters of key DDR (Fig. 2*D*) and cell cycle (*SI Appendix*, Fig. S5*A*) genes were often occupied by HDAC1 and HDAC2 along with active histone marks and transcription machinery (Fig. 2*D*), supporting a role for HDACs in sustaining gene expression through promoter-driven transcriptional activity. Consistently, HDAC1/2-bound promoters were highly enriched for SP1, KLF1/3, NF-Y, and ELK4 motifs (*SI Appendix*, Fig. S6*A*), which are canonical promoter-proximal elements shown to regulate transcription outputs essential for DDR, cell cycle, and other oncogenic processes in cancer (56–59). In contrast, genes associated with intergenic HDAC1/2-occupied sites were strongly upregulated by entinostat (Fig. 2*C*, right) and enriched for cell differentiation, apoptosis, and other HDACi-induced pathways (*SI Appendix*, Figs. S4*B* and S5*B*), implicating a location-dependent repressive function of HDACs at intergenic regions, which are often enhancer-rich. Furthermore, HDAC1/2-bound intergenic sites were enriched for motifs of AP-1 family transcription factors (FRA1, FOS, JUNB, FRA2) and the AP-1-related factor ATF3 (*SI Appendix*, Fig. S6*B*), which are commonly associated with enhancer elements and dynamically regulated in tumorigenesis (60–62). Together, these results suggest dual, genomic location-dependent roles of HDAC1/2: they potentially maintain active transcription at promoters, including at DDR genes, while repressing transcription activity driven by intergenic enhancers.

### HDACi Causes H3K27ac to Accumulate Preferentially at Intergenic Regions Over Promoters

To further dissect the roles of HDACs in transcriptional regulation, we next examined how HDAC inhibition alters the genomic distributions of H3K27ac after 6 h and 24 h of entinostat treatment, timepoints when DDR gene suppression was first established and at a maximum, respectively (Fig. 1*G* and *H*). Consistent with the global increases in H3K27ac and total H3 acetylation (*SI Appendix*, Fig. S7*A* and *B*), entinostat markedly expanded the total number of H3K27ac-occupied sites (3.2-fold at 6 h and 2.5-fold at 24 h; Fig. 3*A*). Although all annotated genomic regions gained peaks after 6 h, intergenic regions showed the greatest expansion (4-fold), followed by genic (3.3-fold) and promoter (2-fold) regions (Fig. 3*A*). As a result, the proportion of peaks that were intergenic increased from 29% to 36% at 6 h, the proportion of promoter peaks decreased from 23% to 15%, and the genic proportion was largely unchanged, with all these proportions roughly persisting at 24 h (Fig. 3*A*). Collectively, these results indicate a preferential enrichment of H3K27ac at intergenic regions over promoters upon HDAC inhibition.

**Fig. 3.**
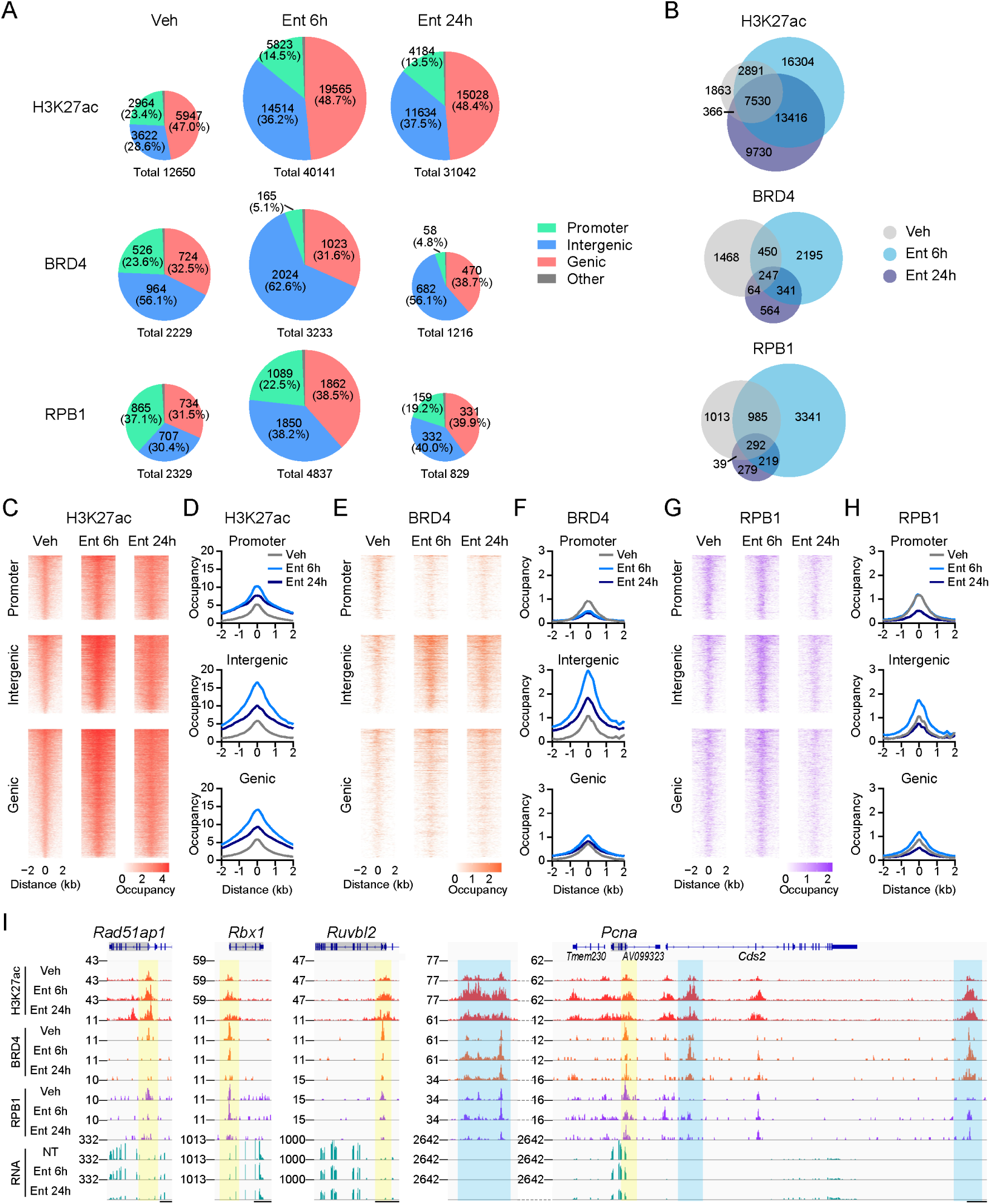
Entinostat preferentially increases intergenic H3K27ac, diverting BRD4 and Pol II from DDR gene promoters. (*A*) Numbers and percentages of annotated H3K27ac, BRD4, and RPB1 peaks in FC1245 cells treated with Veh or Ent (5 µM, 6h or 24h) by CUT&RUN. (*B*) Venn diagrams showing H3K27ac, BRD4, and RPB1 peak distributions across treatments. (*C*–*H*) Heatmaps and average profiles of H3K27ac (*C* and *D*), BRD4 (*E* and *F*), and RPB1 (*G* and *H*) occupancy across treatments at baseline H3K27ac peaks (Veh). (*I*) Genome browser tracks of representative DDR gene loci illustrating Ent-induced changes of H3K27ac, BRD4, and RPB1 at promoters (yellow) and in selected adjacent intergenic/genic regions (blue) along with RNA changes. NT, no treatment. Scale bar, 5 kb.

Pre-existing H3K27ac peaks (as detected in vehicle-treated cells) largely persisted through entinostat treatment (Fig. 3*B* and *SI Appendix*, Fig. S8*A*) and showed region-dependent gains. Intergenic regions displayed the highest gain of H3K27ac, genic regions were intermediate, and promoters were the lowest, despite comparable baseline occupancy across all regions (Fig. 3*C* and *D*). Similar region-dependent H3K27ac increases were also observed at HDAC1/2-bound sites (*SI Appendix*, Fig. S9*A* and *B*), reflecting the widespread co-occupancy of HDAC1/2 and H3K27ac (Fig. 2*B*). In addition, entinostat induced extensive de novo acetylation (29,720 sites at 6 h), mainly in genic (50%) and intergenic (38%) regions, with relatively few at promoters (12%) (Fig. 3*B* and *SI Appendix*, Fig. S8*A*). Among these, intergenic sites showed slightly higher H3K27ac gain compared to genic and promoter sites (*SI Appendix*, Fig. S10*A* and *B*). Together, these results demonstrate that HDAC inhibition disrupts the balanced promoter-intergenic distribution of H3K27ac by preferentially accumulating acetylation at intergenic regions.

### HDACi Diverts Transcriptional Machinery from Active Promoters to Intergenic Regions

Given that histone acetylation recruits the bromodomain-containing reader BRD4, which in turn engages RNA Pol II to drive transcription, we next analyzed how HDACi-induced acetylation change affects genome-wide BRD4 and RPB1 occupancy. After 6 h of entinostat treatment, BRD4 bound to 1.5-fold more genomic sites (Fig. 3*A*) without major change in protein abundance (*SI Appendix*, Fig. S7*C* and *D*), suggesting that the genomic occupancy change mainly results from repartitioning of BRD4 across the genome. Intergenic BRD4 peaks nearly doubled, increasing their share from 56% to 63% of total peaks, whereas promoter-associated peaks dropped more than 3-fold, representing only 5.1% of total peaks after 6 h compared to 24% at baseline (Fig. 3*A*). In total, 1,532 baseline BRD4 peaks were lost (69%; Fig. 3*B*), including 88% of the promoter, 65% of genic, and 61% of intergenic peaks (*SI Appendix*, Fig. S8*B*). Concurrently, 2,536 de novo BRD4 peaks emerged (Fig. 3*B*), mainly at intergenic (65%) and genic (30%) regions, with few at promoters (4%) (*SI Appendix*, Fig. S8*B*), highlighting the redistribution of BRD4 from promoters to intergenic regions. Notably, after 24 h of entinostat treatment there was a significant decrease in BRD4 occupancy across all regions relative to 6 h (Fig. 3*A* and *B*), which corresponded to a modest decrease in total BRD4 protein (*SI Appendix*, Fig. S7*C* and *D*); however, the promoter-to-intergenic redistribution persisted (Fig. 3*A* and *SI Appendix*, Fig. S8*B*).

Upon entinostat treatment, BRD4 was redistributed to pre-existing intergenic sites (Fig. 3*E* and *F* and *SI Appendix*, Fig. S9*C* and *D*), where H3K27ac showed the greatest gains (Fig. 3*C* and *D*, and *SI Appendix*, Fig. S9*A* and *B*). In contrast, its occupancy markedly declined at promoters (Fig. 3*E* and *F* and *SI Appendix*, Fig. S9*C* and *D*), where H3K27ac increased the least (Fig. 3*C* and *D*, and *SI Appendix*, Fig. S9*A* and *B*). Consequently, BRD4 occupancy ranked highest at intergenic, intermediate at genic, and lowest at promoter sites (Fig. 3*E* and *F* and *SI Appendix*, Fig. S9*C* and *D*), paralleling the altered H3K27ac distribution (Fig. 3*C* and *D* and *SI Appendix*, Fig. S9*A* and *B*). De novo acetylated sites followed the same hierarchy in BRD4 occupancy, showing strong BRD4 recruitment to intergenic sites, modest binding at genic, and negligible at promoters (*SI Appendix*, Fig. S10*C* and *D*). After 24 h of entinostat treatment, these patterns largely persisted, with BRD4 occupancy at both pre-existing and de novo intergenic H3K27ac sites decreasing slightly relative to 6 h (Fig. 3*E* and *F* and *SI Appendix*, Fig. S10*C* and *D*). Collectively, these findings indicate that HDAC inhibition rapidly diverts BRD4 from promoters toward intergenic regions, mirroring the acetylation imbalance.

Pol II (RPB1) exhibited similar genomic dynamics as BRD4 under HDACi at 6 h. Entinostat treatment slightly increased RPB1 protein (*SI Appendix*, Fig. S7*C* and *D*) and roughly doubled the number of RPB1 peaks at this time point (Fig. 3*A*). Although all genomic categories gained RPB1 occupancy, the share of promoter peaks dropped from 37% to 22.5%, while the intergenic and genic shares rose from 30% to 38% and 31% to 39%, respectively (Fig. 3*A*). RPB1 occupancy was lost at 45% of baseline sites (1,052 sites; Fig. 3*B*), with the most prominent loss at promoters (50%) compared to genic (42%) and intergenic (42%) regions (*SI Appendix*, Fig. S5*C*). Meanwhile, 3,560 de novo RPB1 peaks were induced, approximately three-fold more than the loss (Fig. 3*B*). The majority of de novo binding occurred at intergenic (40%) and genic (40%) regions, and fewer at promoters (19%; *SI Appendix*, Fig. S8*C*).

Upon entinostat treatment, RPB1 occupancy increased most prominently at intergenic sites, moderately at genic, but not at promoter sites (Fig. 3*G* and *H* and *SI Appendix*, Fig. S9*E* and *F*), largely consistent with the redistributed H3K27ac and BRD4 patterns (Fig. 3*C*–*F* and *SI Appendix*, Fig. S9*A*–*D*). De novo H3K27ac sites showed the same RPB1 occupancy hierarchy across annotations (*SI Appendix*, Fig. S10*E* and *F*). At 24 h of entinostat treatment, total RPB1 peaks fell to a level 2.8-fold below baseline (Fig. 3*A*). While the promoter-to-intergenic redistribution seen at 6 h largely persisted at 24 h (Fig. 3*A* and *SI Appendix*, Fig. S8*C*), RBP1 occupancy was markedly reduced across all regions at 24 h compared to 6 h, showing more pronounced changes than those observed in BRD4 occupancy (Fig. 3*G* and *H* and *SI Appendix*, Fig. S9*E* and *F* and S10*E* and *F*). This agrees with our findings that total RPB1 protein decreases by ∼40% at 24 h of entinostat treatment, compared with only a minor reduction in BRD4 (*SI Appendix*, Fig. S7*C* and *D*).

Together, the genomic analyses reveal that entinostat induces H3K27ac accumulation preferentially at intergenic regions, rapidly diverting BRD4 and Pol II away from promoters to intergenic loci and thereby attenuating promoter-driven transcription activity. These dynamics were frequently evident at promoters of entinostat-downregulated genes, including those regulating DDR (Fig. 3*I*) and cell cycle (*SI Appendix*, Fig. S11*A*). These promoters showed reduced BRD4 and RPB1 occupancy, despite modest H3K27ac gains (Fig. 3*I* and *SI Appendix*, Fig. S11*A*, highlighted in yellow). In contrast, intergenic and genic regions with greater H3K27ac gains accumulated both BRD4 and RPB1 (Fig. 3*I*, exemplified by *Pcna*-adjacent loci highlighted in blue). These dynamics were frequently observed in genomic regions near entinostat-upregulated genes (*SI Appendix*, Fig. S11*B*), suggesting that entinostat may induce their transcriptional activation through increased activity of intergenic enhancers. Overall, we conclude that HDAC inhibition induces intergenic-biased H3K27ac accumulation, which redistributes BRD4 and Pol II from active promoters to intergenic elements and leads to suppression of promoter-driven transcription, including at DDR genes.

### HDACi Enhances the Efficacy of DNA-Damaging and DDR-Targeting Agents in PDAC models

Recognizing that HDACi-mediated suppression of DDR genes may sensitize PDAC cells to genotoxic stress, we next tested whether HDAC inhibition enhances the efficacy of DNA-damaging agents currently used in PDAC therapeutic regimens, like platinum-based crosslinkers. Indeed, combining entinostat with cisplatin or oxaliplatin increased DNA damage, as indicated by elevated γ-H2A.X (Fig. 4*A* and *B*), and reduced cell viability in a concentration-dependent (*SI Appendix*, Fig. S12*A* and *B*) and synergistic manner (Fig. 4*C*–*F*) across multiple PDAC cell lines.

**Fig. 4.**
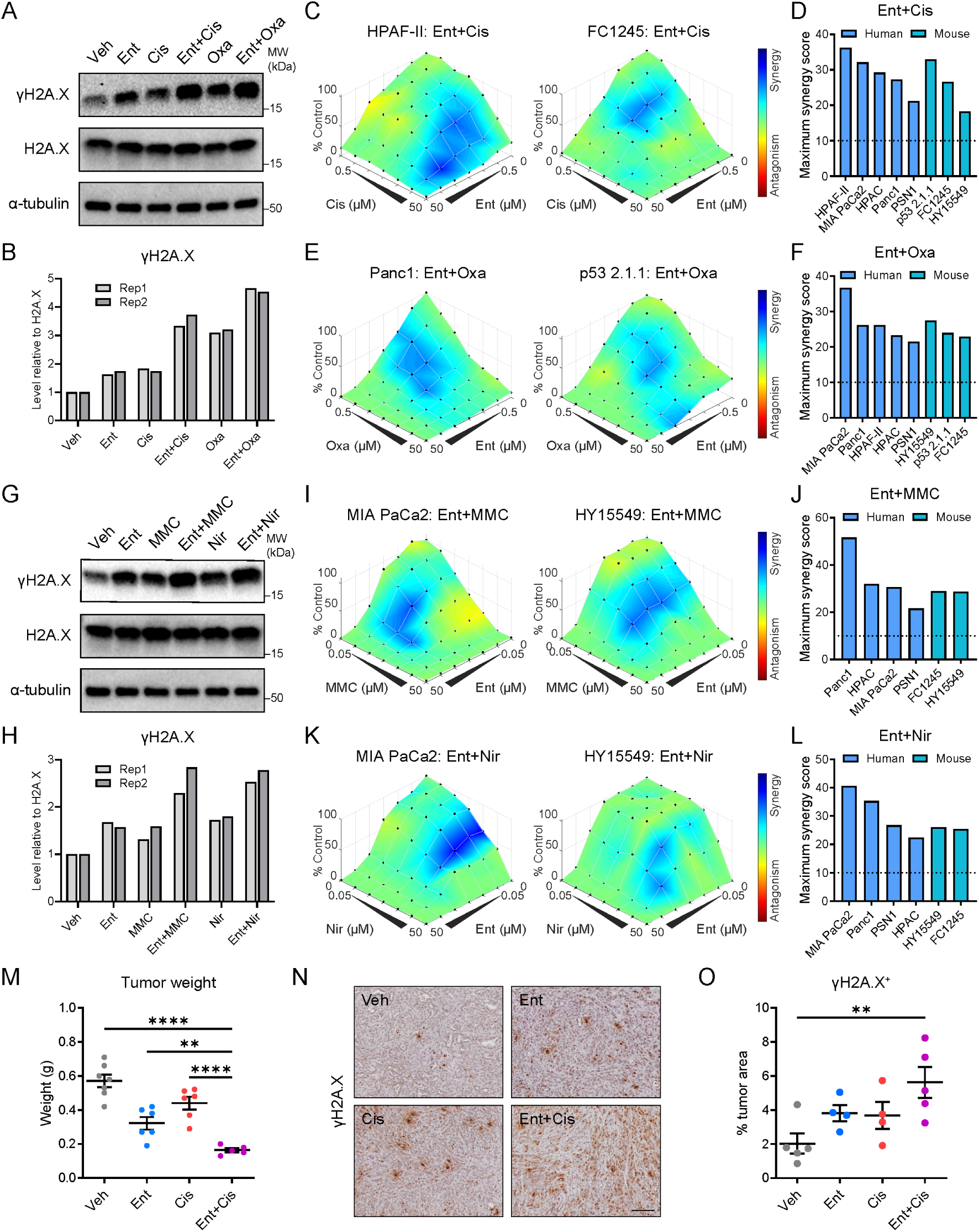
Entinostat enhances the efficacy of DNA-damaging and DDR-targeting agents in PDAC. (*A* and *B*) Representative Western blots (*A*) and quantification (*B*) showing increased γH2A.X (normalized to H2A.X) in FC1245 cells after 6 h of Ent treatment (1 µM) combined with cisplatin (Cis, 1 µM) or oxaliplatin (Oxa, 5 µM). (*C*–*F*) Representative plots and maximum scores (highest single agent model) from synergy analysis on viability of human and mouse PDAC cells under graded co-treatments of Ent with Cis (*C* and *D*) or Oxa (*E* and *F*). (*G* and *H*) Representative Western blots (*G*) and quantifications (*H*) showing increased γH2A.X in FC1245 after 6 h combined treatments of Ent (1 µM) with mitomycin C (MMC, 0.5 µM) or niraparib, (Nir, 2 µM). (*I*–*L*) Synergy plots and maximum scores from co-treatments of Ent and MMC (*I* and *J*) or Nir (*K* and *L*) in PDAC cells. Western blotting, *n* = 2 independent experiments; synergy analysis, *n* = 3 cell sample replicates. (*M*) Tumor weights from orthotopic transplantation models (FC1245) treated for two weeks with Ent (20 mg/kg, daily) and/or Cis (0.5 mg/kg, q3d). *n* = 7 (Veh), 6 (Ent, Cis), or 5 (Ent+Cis) mice. (*N* and *O*) Representative IHC images and quantifications of γH2A.X in tumors across treatment groups. *n* = 5 (Veh, Ent+Cis) or 4 (Ent, Cis) tumors. Scale bar, 50 µm. Data are presented as mean values ± SEM (*M* and *O*). **, *p <* 0.01; ****, *p* < 0.0001. One-way ANOVA.

Since oxaliplatin is used clinically in the FOLFIRINOX regimen, which also contains irinotecan (a prodrug of the topoisomerase I inhibitor SN38) and 5-FU (a pyrimidine base analog), we further evaluated whether these drugs influence the synergy between entinostat and oxaliplatin. Notably, entinostat and oxaliplatin exhibited enhanced synergy in the combined presence of SN38 and 5-FU (*SI Appendix*, Fig. S13*A*). SN38 and 5-FU also each individually synergized with entinostat (*SI Appendix*, Fig. S13*B*), indicating the ability of HDAC inhibition to exacerbate cellular stress induced by diverse DNA-damaging mechanisms.

Extending our analysis to additional classes of DNA-damaging and DDR-targeting agents, we found that entinostat synergizes with representative drugs from multiple classes of these therapeutics (*SI Appendix*, Table S1), including mitomycin C (a non-platinum DNA crosslinker) and niraparib (a PARP1/2 inhibitor), with enhanced cytotoxicity observed in multiple PDAC cell lines (Fig. 4*I*–*L* and *SI Appendix*, Fig. S12*C* and *D*). Synergy was also observed when entinostat was combined with the ATM inhibitor AZD0156 and the DNA-intercalating agent doxorubicin (*SI Appendix*, Fig. S13*C*).

To evaluate the in vivo efficacy of combining HDACi with DNA-damaging agents, we treated syngeneic orthotopic PDAC models with entinostat and cisplatin. The combination treatment significantly reduced tumor burden compared to either agent alone (Fig. 4*M*). This was paralleled by a corresponding increase in intratumoral DNA damage, as measured by γH2A.X (Fig. 4*N* and *O*). Together, these findings demonstrate that HDAC inhibition broadly enhances the efficacy of DNA damage-inducing agents, including clinically relevant regimens, establishing the potential of HDACi-based drug combinations for the treatment of PDAC.

### Tumor-Selective Delivery of HDACi by BPD Nanoparticles Enhances Its Therapeutic Potential

The clinical application of HDACis has been limited by their narrow or often absent therapeutic windows (45, 46). To overcome this challenge, we tested if a BPD nanoparticle platform could achieve tumor-selective delivery of entinostat, thereby minimizing systemic HDAC inhibition and related toxicity (48, 49). Using a fluorescence-labeled BPD probe (63), we confirmed that BPDs selectively accumulate in orthotopic PDAC tumors compared to adjacent normal pancreas and other organs (Fig. 5*A* and *B*), consistent with the tumor-enriched biodistribution previously observed for BPDs in other cancer models (48, 49).

**Fig. 5.**
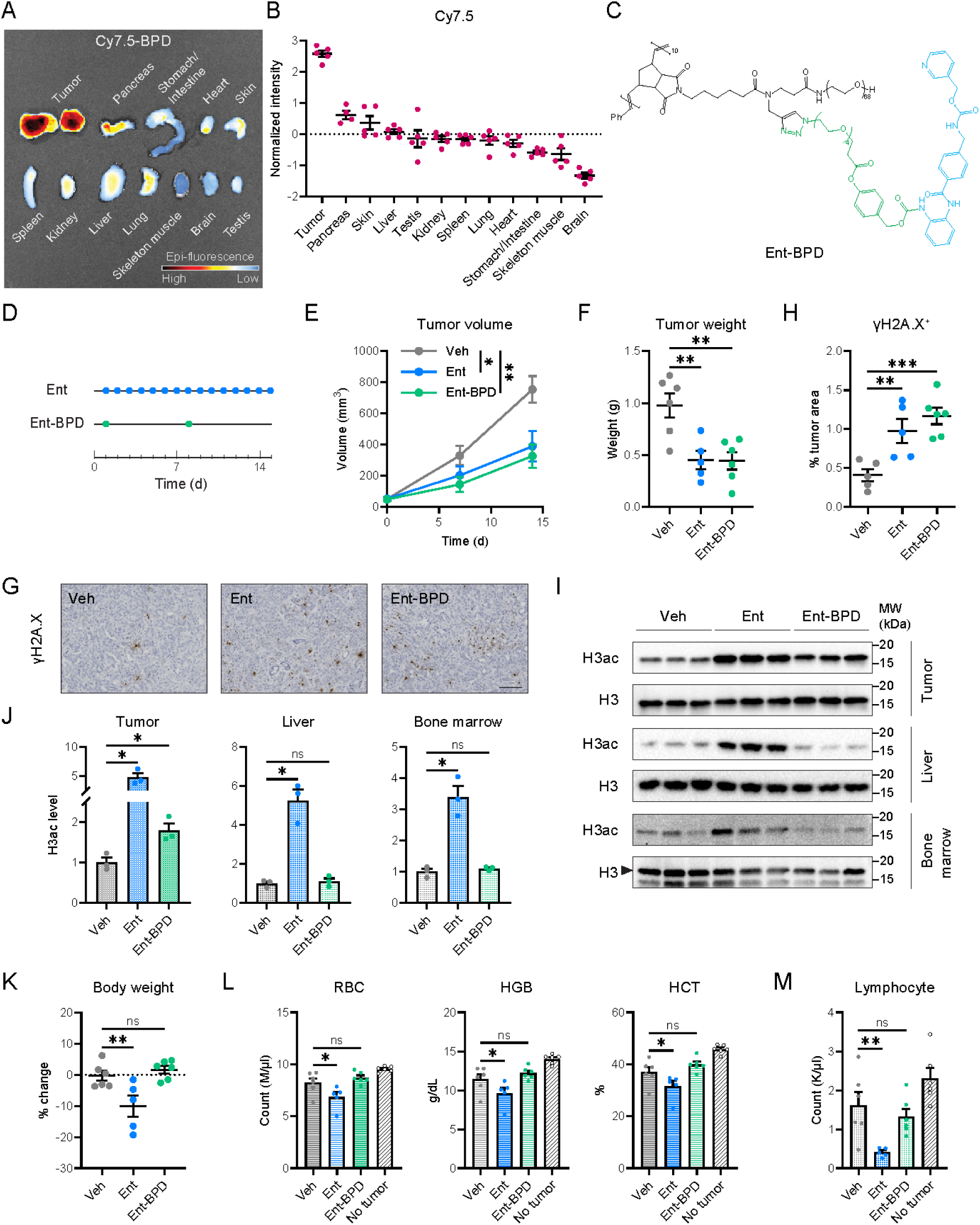
In vivo tumor selectivity, efficacy, and enhanced safety of Ent-BPD. (*A* and *B*) Representative image (*A*) and quantification (*B*) from fluorescence imaging of tumors and organs from HY15549 orthotopic models administrated Cy7.5-BPD (250 mg/kg body weight) 24 h before sacrifice. *n* = 5 mice (one mouse lacking sufficient adjacent normal pancreas). (*C*) Chemical structure of Ent-BPD highlighting the Ent moiety (blue) and esterase-cleavable linker (green). (*D*) Two-week treatment schemes of Ent (15 mg/kg, daily) or Ent-BPD (15 mg FD eq/kg, q7d) in HY15549 orthotopic models enrolled by ultrasound imaging at day 0, treated from day 1, and sacrificed at day 15. FD eq, free drug equivalent. (*E* and *F*) Tumor volumes by longitudinal ultrasound (*E*) and tumor weights at endpoint (*F*) over 2-week treatments. *n* = 6 (Veh, Ent-BPD) or 5 (Ent) mice. (*G* and *H*) Representative IHC images (*G*) and quantification (*H*) of γH2A.X in tumors across treatment groups. *n* = 6 (Ent-BPD) or 5 (Veh, Ent) tumors. Scale bar, 50 µm. (*I* and *J*) Western blots (*I*) and quantifications (*J*) of H3ac levels (relative to H3) in tumor, liver, and bone marrow samples at endpoint, normalized to Veh. *n* = 3 mice. (*K*) Body weight changes after 1 week of treatment relative to pre-treatment. (*L* and *M*) Red blood cell (RBC) count, hemoglobin (HGB) concentration, hematocrit (HCT) percentage (*L*), and lymphocyte count (*M*) in treated tumor-bearing mice and non-tumor-bearing mice at endpoint. *n* = 6 (Veh, Ent-BPD, No tumor) or 5 (Ent) mice. Data are presented as mean values ± SEM. *, *p* < 0.05; **, *p <* 0.01; ***, *p* < 0.001; ns, not significant. Two-way ANOVA (*E*, day 14); one-way ANOVA (*F*, *H*, and *J*–*M*).

We next synthesized entinostat-loaded BPD (Ent-BPD; Fig. 5*C* and *SI Appendix*, Supporting Methods). Leveraging the functionalizable amine group, we conjugated entinostat to the polynorbornene-based backbone through a phenyl ester linker that, upon hydrolysis and 1,6-elimination, enables controlled, tumor-selective release of unmodified entinostat (48, 64). Specifically, a reactive linker (Compound 1; *SI Appendix*, Supporting Methods) was coupled to entinostat via carbamate formation (Compound 2). This azide-containing prodrug was then appended onto an alkyne-containing macromonomer (56) via copper-catalyzed alkyne-azide cycloaddition (CuAAC), resulting in entinostat-loaded macromonomer (Ent-M). These compounds were analyzed using ^1^H and ^13^C nuclear magnetic resonance spectroscopy (NMR), mass spectrometry (MS), and matrix-assisted laser desorption ionization time-of-flight mass spectrometry (MALDI-MS) where appropriate (*SI Appendix*, Figs. S14–17). Ring-opening metathesis polymerization (ROMP) of Ent-M resulted in Ent-BPD with a number-averaged degree of polymerizations of 10. Ent-BPD was characterized by gel permeation chromatography (GPC), which revealed near-quantitative conversion (>95%; *SI Appendix*, Fig. S18), and by dynamic light scattering (DLS), which suggested a hydrodynamic diameter (D_h_) of ∼10 nm (*SI Appendix*, Fig. S19), a size range shown to achieve efficient tumor accumulation.

Compared to daily free entinostat dosing at 15 mg/kg, weekly Ent-BPD treatment at a free-drug equivalent dose of 15 mg/kg—one-seventh of the cumulative dose in the free entinostat regimen (Fig. 5*D*)—achieved similar tumor growth inhibition (Fig. 5*E* and *F*) and comparable γH2A.X accumulation (Fig. 5*G* and *H*), suggesting that BPD can maintain entinostat therapeutic efficacy while reducing systemic exposure. Indeed, while free entinostat enhanced H3 acetylation broadly across tumors and normal tissues (e.g., liver and bone marrow), Ent-BPD limited H3 acetylation increase to tumors (Fig. 5*I* and *J*), demonstrating tumor-selective HDAC inhibition.

Importantly, Ent-BPD treatment avoided the systemic side effects associated with free entinostat, including body weight loss (Fig. 5*K*), anemia (manifested by reduced red blood cell counts, hemoglobin, and hematocrit; Fig. 5*L*), and lymphopenia (Fig. 5*M*). The ability to achieve anti-tumor efficacy with minimum systemic exposure and toxicity highlights the therapeutic potential of BPD-mediated HDACi delivery.

We next tested whether entinostat-BPD can enhance the efficacy of DNA-damaging agents shown to synergize with HDACi (Fig. 4). In combination with oxaliplatin, Ent-BPD significantly reduced the growth of orthotopic PDAC tumors in mice, compared to either agent alone (Fig. 6*A* and *B*). This enhanced efficacy was achieved without body weight loss (Fig. 6*C*), supporting Ent-BPD as an effective and tolerated addition to current PDAC therapeutic regimens.

**Fig. 6.**
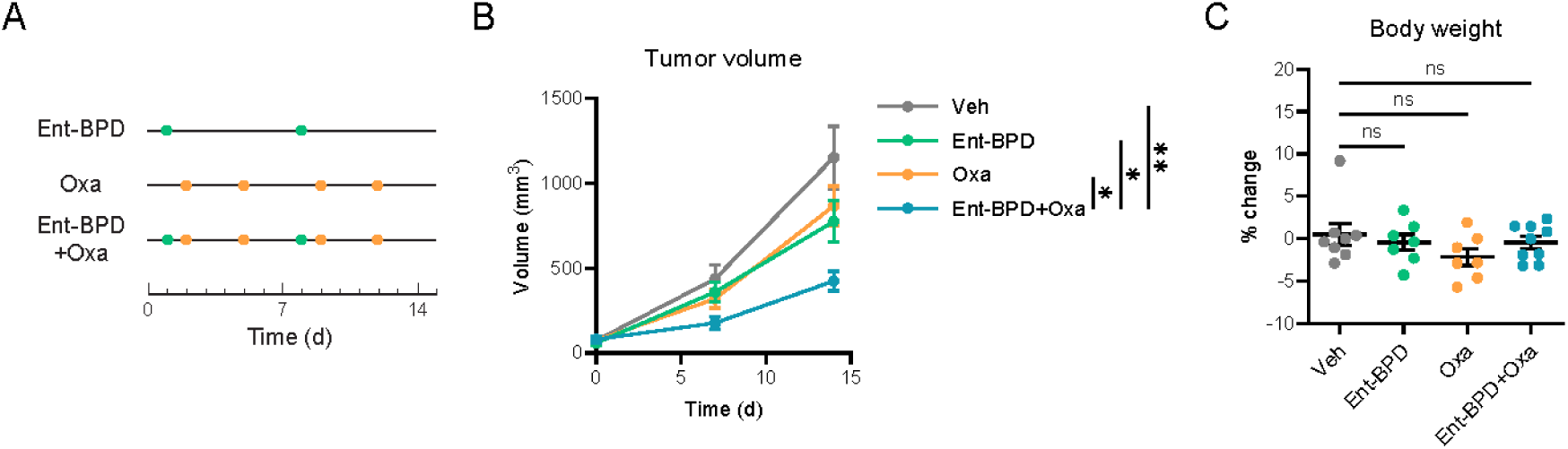
Ent-BPD enhances the efficacy of DNA-damaging agents. (*A*) Two-week treatment schemes of Ent-BPD (5 mg FD eq/kg, q7d), Oxa (0.5 mg/kg, twice weekly), or the combination in HY15549 orthotopic models. (*B* and *C*) Tumor volumes by longitudinal ultrasound over 2 weeks (*B*) and body weight changes after 1 week (*C*) of Ent-BPD and/or Oxa treatments. *n* = 7 (Ent-BPD, Oxa), 8 (Veh), 9 (Ent-BPD+Oxa) mice. Data are presented as mean values ± SEM. *, *p* < 0.05; **, *p <* 0.01; ns, not significant. Two-way ANOVA (*B*, day 14); one-way ANOVA (*C*).

## DISCUSSION

Our study identifies a critical role of Class I HDACs in sustaining the DDR in PDAC and elucidates how HDACs facilitate DDR gene expression. We show that HDAC1/2 maintain a proper distribution of histone acetylation between promoters and intergenic regions, supporting BRD4 and Pol II recruitment to DDR gene promoters and enabling promoter-driven transcription (schematic model in Fig. 7). Importantly, while HDAC inhibition increases histone acetylation globally, it does so unevenly—driving the preferential accumulation of acetylation at intergenic regions over promoters. This imbalanced acetylation distribution diverts BRD4 and Pol II away from active promoters to intergenic loci, attenuating promoter-driven transcription activity at DDR genes (Fig. 7).

**Fig. 7.**
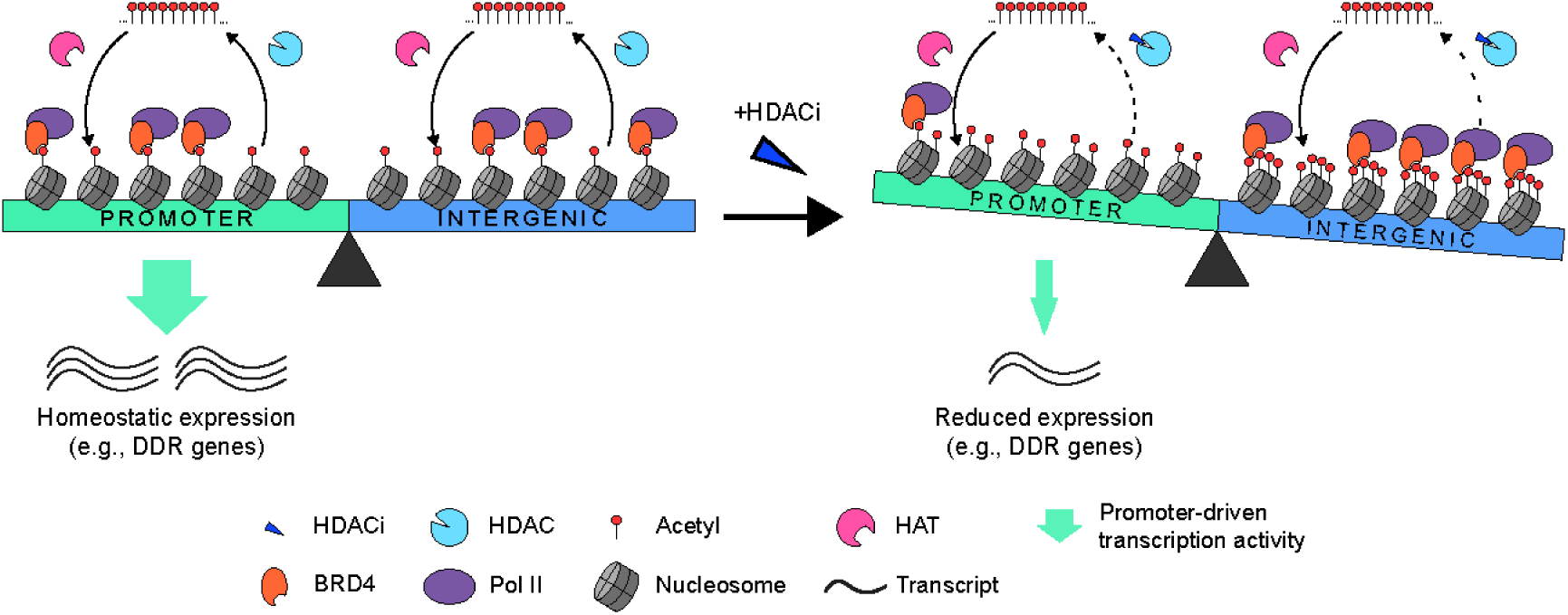
Schematic model illustrating how HDACs regulate DDR gene expression by maintaining balanced genome-wide distribution of promoter and intergenic acetylation. Under the baseline condition, DDR genes are highly expressed in PDAC cells due to promoter-driven transcription activity, with balanced distribution of histone acetylation between promoters and intergenic regions through coordinated actions of histone acetyltransferases (HATs) and HDACs. HDAC inhibition disrupts this balance by preferentially increasing acetylation at intergenic regions, leading to redistribution of BRD4 and Pol II from promoters to intergenic regions and consequent attenuation of promoter-driven transcription, including that of DDR genes.

These findings add to accumulating evidence that HDACs sustain active transcription at promoters (30–32, 34–36) and regulate the genomic dynamics of BRD4 and Pol II (32, 36, 65, 66). HDAC activity has also been linked to enhancers, including those frequently located in intergenic regions (34, 67). In line with this, we find that intergenic HDAC1/2 occupancy correlates with gene upregulation (Fig. 2*C*), greater H3K27ac gain, and increased BRD4 and Pol II occupancy upon HDAC inhibition (Figs. 3*C*–*H* and 7). Collectively, these results suggest that HDACs function in a genomic location-dependent manner and may coordinate between promoter- and enhancer-driven transcription programs. Future work, including Hi-C and promoter capture assays to map enhancer-gene interactions, is needed to further define the mechanistic role of HDACs in enhancer-mediated transcription in PDAC.

Mechanistically, the molecular determinants underlying the dual, context-specific activities of HDACs at promoters and intergenic regions remain to be resolved. While we find that HDACs are associated with distinct transcription factor motifs in a location-dependent manner (*SI Appendix*, Fig. S6), average HDAC1/2 occupancy is largely similar between promoters and intergenic regions (*SI Appendix*, Fig. S3). These observations suggest that additional factors contribute to the context-dependent regulation, including differential compositions, dynamics, and catalytic activities of HDAC-containing complexes (52, 68), interplay between HDACs and histone acetyltransferases (30, 34), and contributions from core transcription factors (33, 66).

Interestingly, while the epigenomic redistributions and the transcriptional changes at DDR genes occur rapidly following HDACi treatment (within 6 h; Figs. 1 and 3), we find prolonged HDACi exposure (24 h) sustains or further reinforces DDR gene suppression (Fig. 1). This sustained repression is likely strengthened by additional mechanisms, including reductions in protein translation capacity (Fig. 2*B* and *E*) as well as decreases in the transcriptional machinery (*SI Appendix*, Fig. S7*C* and *D*) and its genomic occupancy (Fig. 3 and *SI Appendix*, Figs. S8 and S10). The mechanistic details and contributions of these later-phase HDACi-induced events remain to be elucidated.

Our study further demonstrates that HDACi-induced suppression of DDR genes creates a therapeutic vulnerability that can be exploited by DNA-damaging agents. Consistent with our findings, downregulation of DNA repair hallmarks under HDACi was also observed in PDAC patient samples from a recent Phase III clinical trial combining entinostat with immune checkpoint blockade (44), underscoring the clinical relevance of our findings. Given the reliance of frontline PDAC chemotherapies on DNA-damaging agents, the integration of clinically advanced HDACi, such as entinostat, into these regimens represents a promising and readily testable therapeutic strategy. To maximize clinical benefit, future studies should incorporate stratification based on DDR competency, including mutational status of DDR components (12). In addition, it will be important to understand how recently developed RAS inhibitors, which are being tested in PDAC patients, impact the observed synergy of combined entinostat and chemotherapy (69).

Our development of BPD-mediated, tumor-selective HDACi delivery achieves potent anti-tumor efficacy with minimal systemic toxicity, thereby improving the therapeutic potential of HDACi. The ability of Ent-BPD to selectively inhibit HDACs within PDAC tumors, together with the versatility of the BPD platform to load diverse therapeutic agents, highlights the potential adaptability of this platform to other PDAC treatment regimens. Moreover, beyond the single-drug BPDs evaluated here, dual-drug BPDs could be engineered to deliver and fine-tune synergistic combinations (49), while antibody-BPD conjugate may further refine tumor specificity (70). Notably, our study focuses on modeling the effects of entinostat-based therapeutics in primary tumors, and the impacts of these therapies on metastatic progression and metastatic tumor growth remain to be determined.

In summary, our study identifies a critical role of Class I HDACs in supporting DDR by maintaining a proper genomic acetylation balance necessary for active transcription, establishes HDAC inhibition as an effective combinatorial strategy to target DDR in PDAC, and introduces a nanoparticle-based delivery approach to exploit this HDAC-mediated dependency. These findings expand our understanding of HDAC function and have the potential to inform the development of new treatment strategies that can significantly improve PDAC patient outcomes. Moreover, given the central role of the DDR in other cancer types and the frequent use of DNA-damaging agents in cancer therapies, this work is likely to have therapeutic relevance across a broad range of cancer types.

## MATERIALS AND METHODS

### Cell Culture, Manipulation, and Viability Assay

In vitro culture, drug treatment, shRNA-mediated depletion, and viability assays using mouse and human PDAC cell lines are described in *SI Appendix*, Materials and Methods.

### RNA-seq, CUT&RUN, and Data Availability

Sample preparation, library construction, and sequencing data analyses for RNA-seq and CUT&RUN are described in *SI Appendix*, Materials and Methods. Sequencing data are deposited in the NCBI Sequence Read Archive (SRA) database under accession number PRJNA1345093, except for RNA-seq data from MIA PaCa2 and FC1245 cells treated with entinostat for 24 h, which are available under PRJNA524175 (38).

### Western Blotting

Protein extraction and Western blotting are described in *SI Appendix*, Materials and Methods.

### Animal Procedures and Ex Vivo Analyses

All animal procedures were approved by the Institute of Animal Care and Use Committee (IACUC) at the Salk Institute. Mouse housing, orthotopic transplantation, drug administration, whole blood collection and analysis, ex vivo fluorescent imaging, and immunohistochemistry (IHC) analysis are described in *SI Appendix*, Materials and Methods.

### BPD Synthesis

Synthesis of entinostat-BPD are described in *SI Appendix*, Materials and Methods with characterization data (*SI Appendix*, Figs. S14–S19). Cy7.5-BPD was synthesized as previously described (63).

### Statistics

Statistical analyses were performed using Prism software (GraphPad, v10.0.2) or provided by the applied databases or algorithms. *p* values are represented by * (*p* < 0.05), ** (*p* < 0.01), *** (*p* < 0.001), or **** (*p* < 0.0001).

## ACKNOWLEDGEMENTS AND FUNDING SOURCES

We thank D. Tuveson, E. Collison, and H. Ying for PDAC cell lines, X. Zhang for biostatistics consultation, and L. Ong and C. Brondos for administrative assistance. This work was funded by grants from the Lustgarten Foundation (Distinguished Scholar Award to R.M.E. and 122215393), the Don and Lorraine Freeberg Foundation, the David C. Copley Foundation, the Wasily Family Foundation, the Paul M. Angell Family Foundation, the NOMIS Foundation (Science of Health), and NIH grants (CA220468, CA014195, and CA265762). G.L. was supported by an NIH National Research Service Award (F32CA217033). This work was supported by the Salk Institute In Vivo Scientific Services and Advanced Biophotonics Cores, the latter funded by NIH-NCI (P30 014195), NIH-NIA (P30 AG068635), the Henry L. Guenther Foundation, and the Waitt Foundation. R.M.E. holds the March of Dimes Chair in Molecular and Developmental Biology at the Salk Institute.

## Author Contributions

G.L., H.V.-T.N., M.D., J.A.J., and R.M.E. designed research. G.L., H.V.-T.N., H.T., H.Z., D.Y.C., G.E., D.C.N., and Y.D. performed research. H.V.-T.N., H.T., H.Z., A.L., and J.A.J. contributed new reagents. G.L., J.Z., H.T., T.G.O., C.L., and R.T.Y. analyzed data. G.L., H.V.-T.N., J.A.J., and R.M.E. wrote the manuscript. T.H., D.E., R.S., A.L., W.F., M.L.T., A.R.A., and M.D. edited the manuscript.

## Classification

1) Biological Sciences: Medical Sciences; 2) Physical Sciences: Chemistry.

