## Supplementary Information for "HDAC Inhibition Sensitizes Pancreatic Tumors to DNA Damage by Global Redistribution of the Transcriptional Machinery"

This file includes:

Figures S1 to S19
Table S1
Materials and Methods
SI Reference

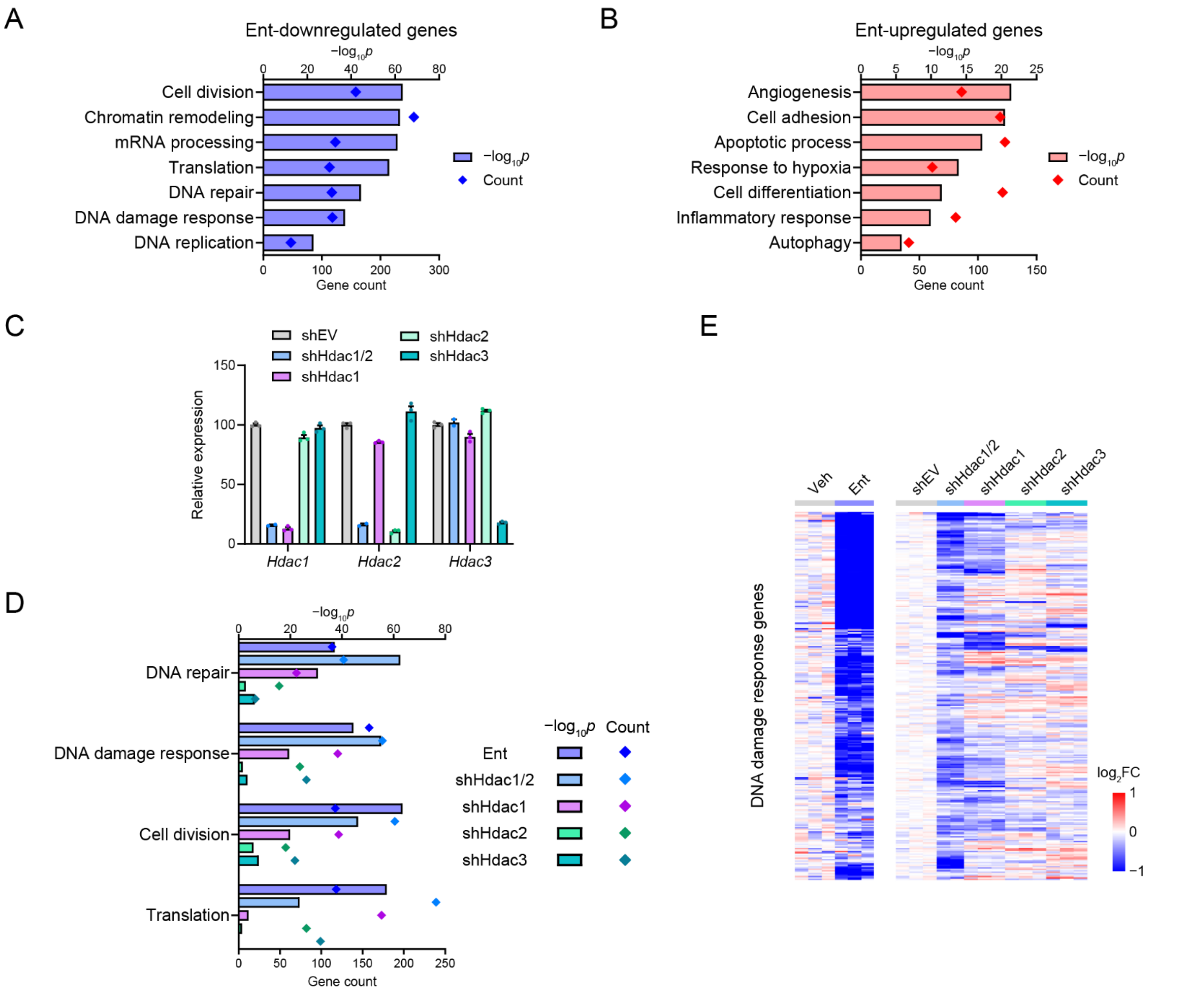


**Fig. S1.** Effects of entinostat treatment and shRNA-mediated HDAC depletions on transcriptional programs in PDAC cells. (*A* and *B*) Representative GO terms enriched in genes significantly downregulated (*A*) and upregulated (*B*) in FC1245 cells treated with Ent (5 µM, 24 h). $n = 2$ independent cell samples. (*C*) RNA-seq based expression data of Class I HDAC genes in FC1245 cells transduced with shRNAs (sh) targeting individual HDACs or both HDAC1 and 2 (shHdac1/2), relative to empty vector (EV) control. Data are presented as mean values ± SEM. (*D*) Enrichment of selected GO terms upon HDAC depletions, compared with Ent treatment. (*E*) Heatmaps showing expression of DDR genes downregulated by both Ent and shHdac1/2 in HDAC-depleted cells and Ent-treated cells. $n = 2$ (shHdac1/2) or 3 (others) independent cell samples.

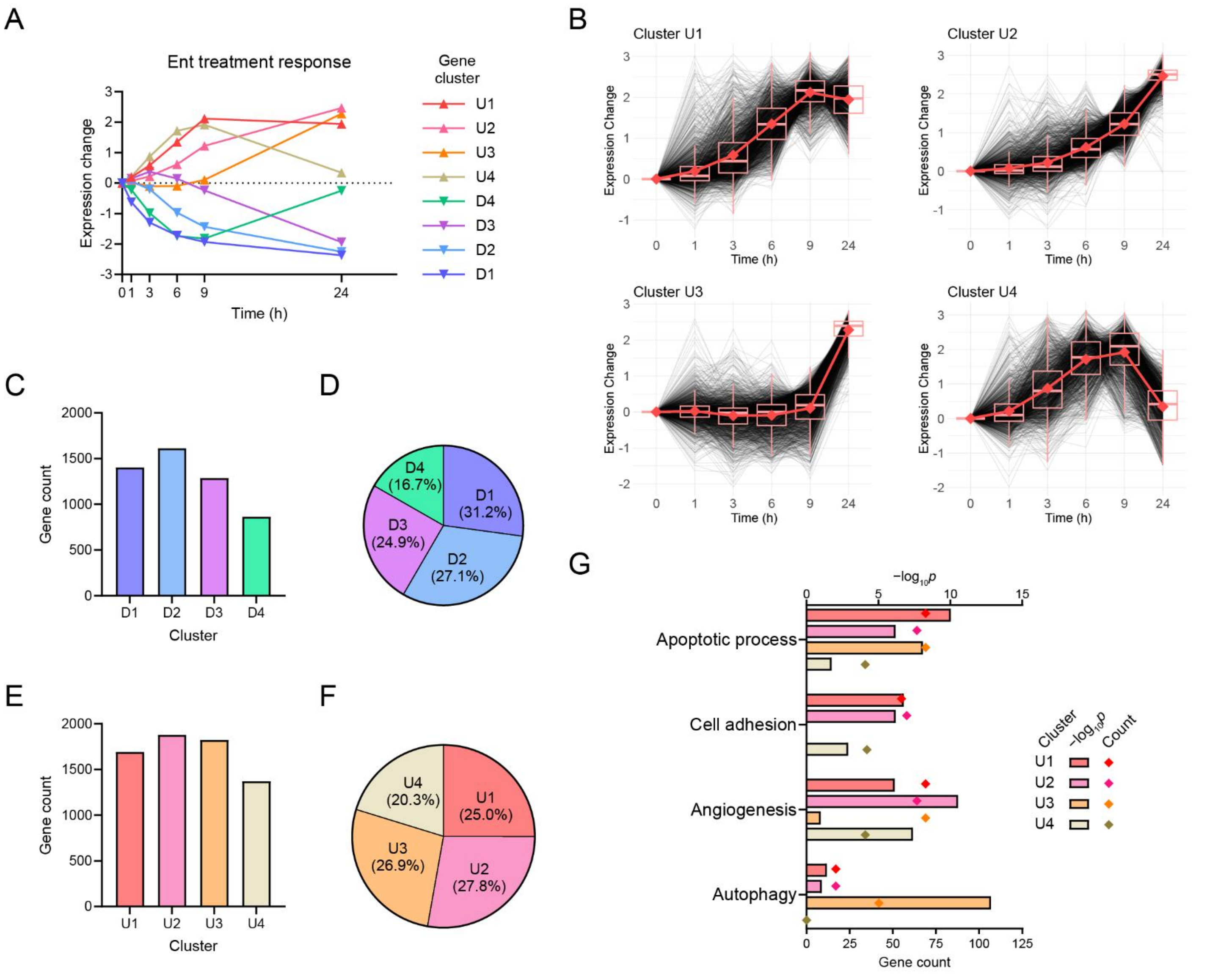


**Fig. S2.** Expression dynamics upon HDACi treatment in PDAC cells. (*A*) Average expression changes of 8 gene clusters identified by temporal gene expression analysis during 24 h treatment of Ent (5 μM) in FC1245, including 4 clusters with upregulation trends (U1–U4) and 4 with downregulation trends (D1–D4). (*B*) Expression dynamics of clusters U1–U4. (*C*–*F*) Gene counts and percentages of individual downregulated (*C* and *D*) and upregulated (*E* and *F*) clusters. (*G*) Enrichment of selected GO terms in upregulated clusters. $n = 3$ independent cell samples.

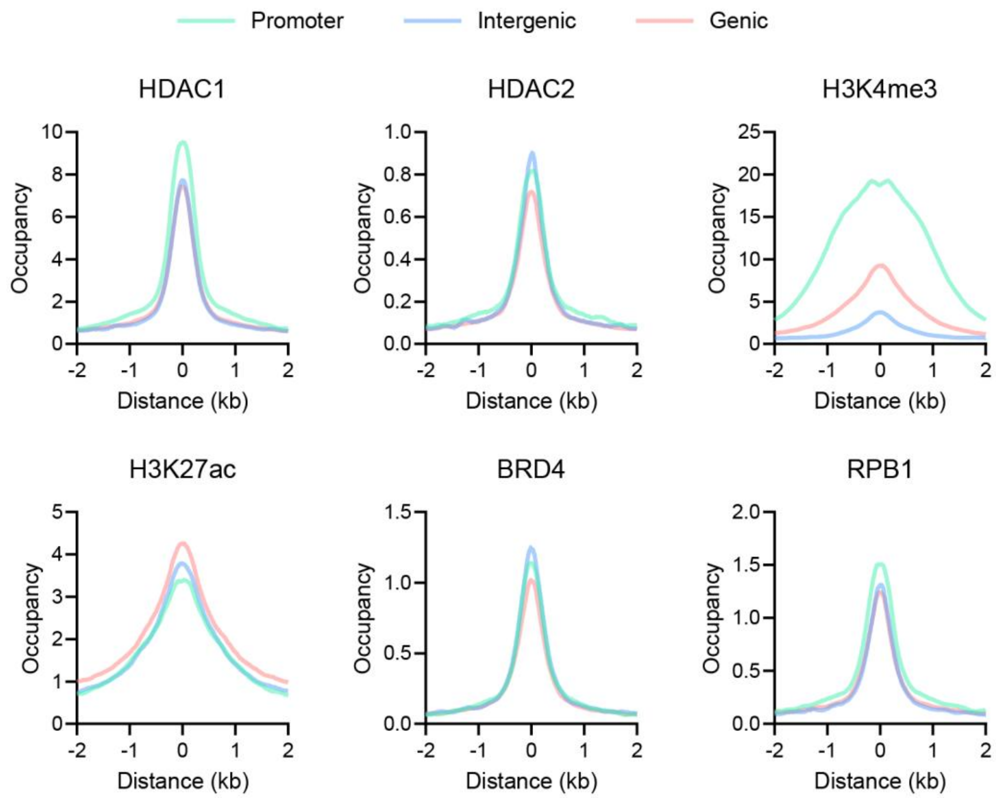


**Fig. S3.** Occupancy of HDAC1, HDAC2, H3K4me3, H3K27ac, BRD4 and RPB1 across HDAC1/2-bound promoter, intergenic, and genic regions.

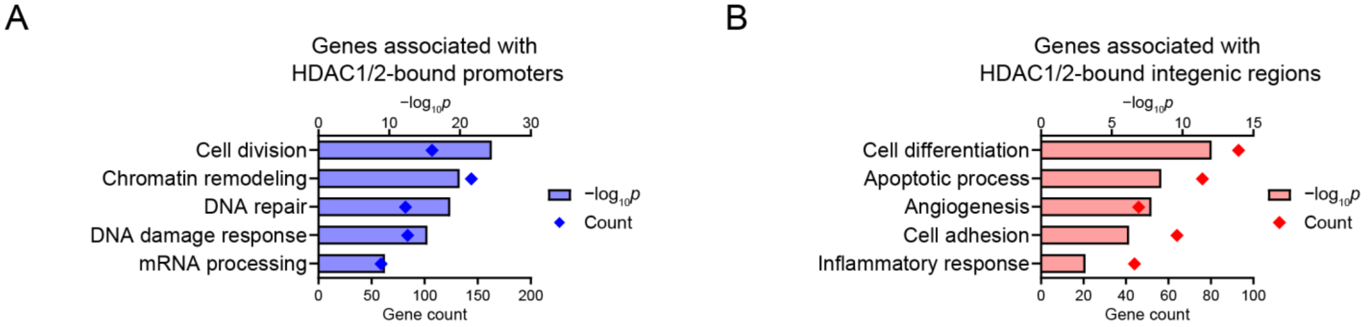


**Fig. S4.** Ontology terms enriched in genes associated with HDAC1/2-bound genomic sites. (*A* and *B*) Representative terms enriched in genes linked to the top 3,000 HDAC1/2-bound promoters (*A*) and intergenic regions (*B*).

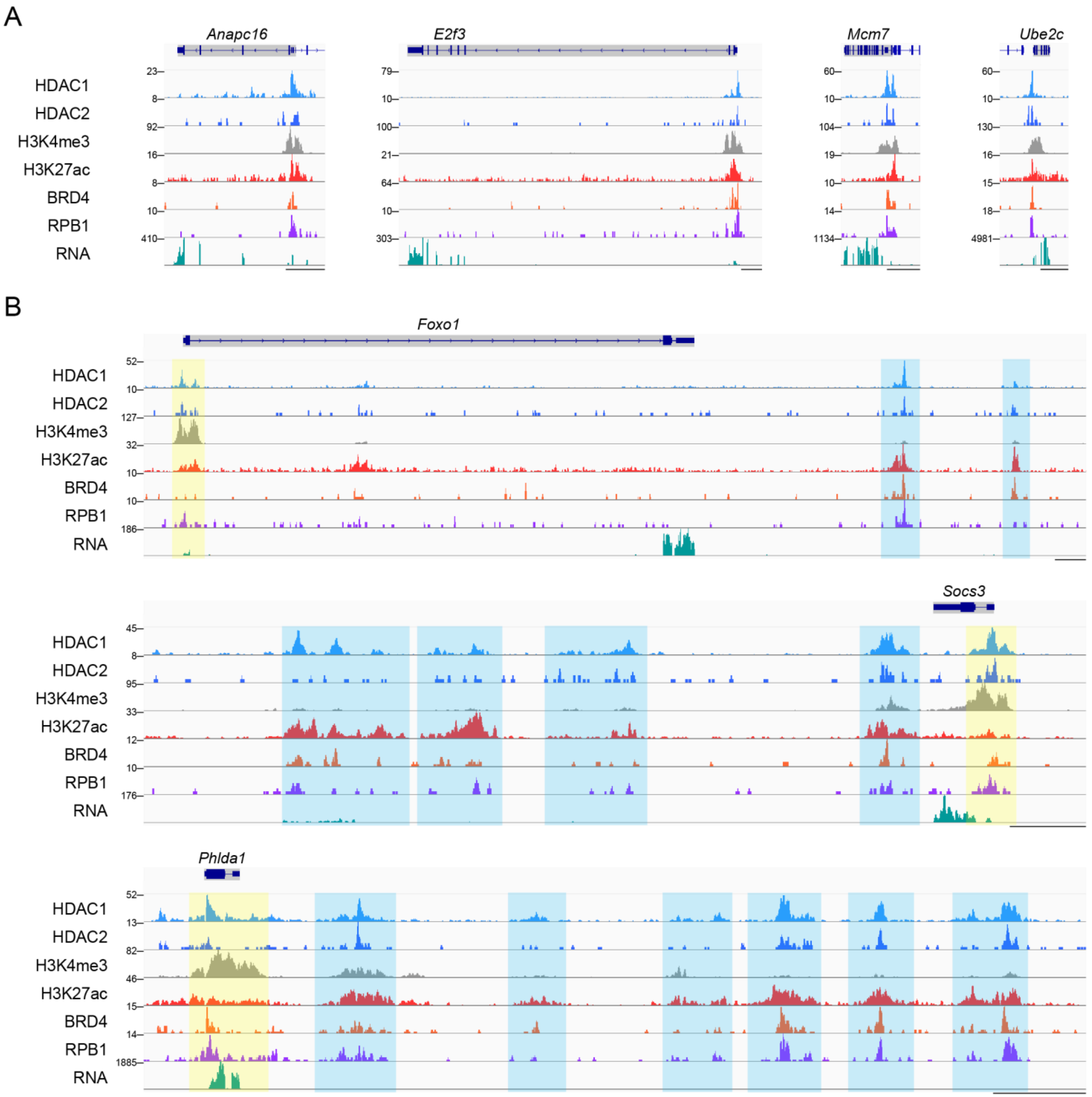


**Fig. S5.** HDAC1/2 occupancy and transcriptional machinery localization at entinostat-regulated gene loci. (*A*) Genome browser tracks showing promoter occupancy of HDAC1/2 and transcriptional machinery at representative entinostat-downregulated cell cycle genes. (*B*) Genome browser tracks showing occupancy of HDAC1/2 and transcriptional machinery at promoters (yellow) and intergenic regions (blue) at representative entinostat-upregulated genes associated with differentiation (*Foxo1*, *Socs3*) and/or apoptosis (*Foxo1*, *Phlda1*). Corresponding RNA output is also shown (*A* and *B*). Scale bar, 5 kb.

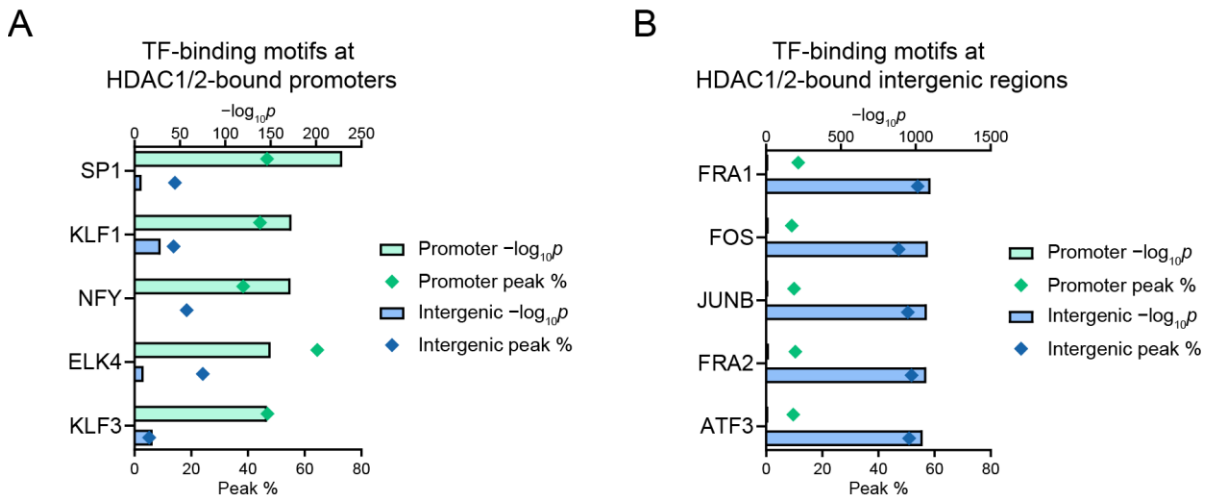


**Fig. S6.** Enriched transcription factor (TF)-binding motifs at HDAC1/2-bound genomic sites. (*A* and *B*) Top 5 motifs for HDAC1/2-bound promoters (*A*) and intergenic regions (*B*).

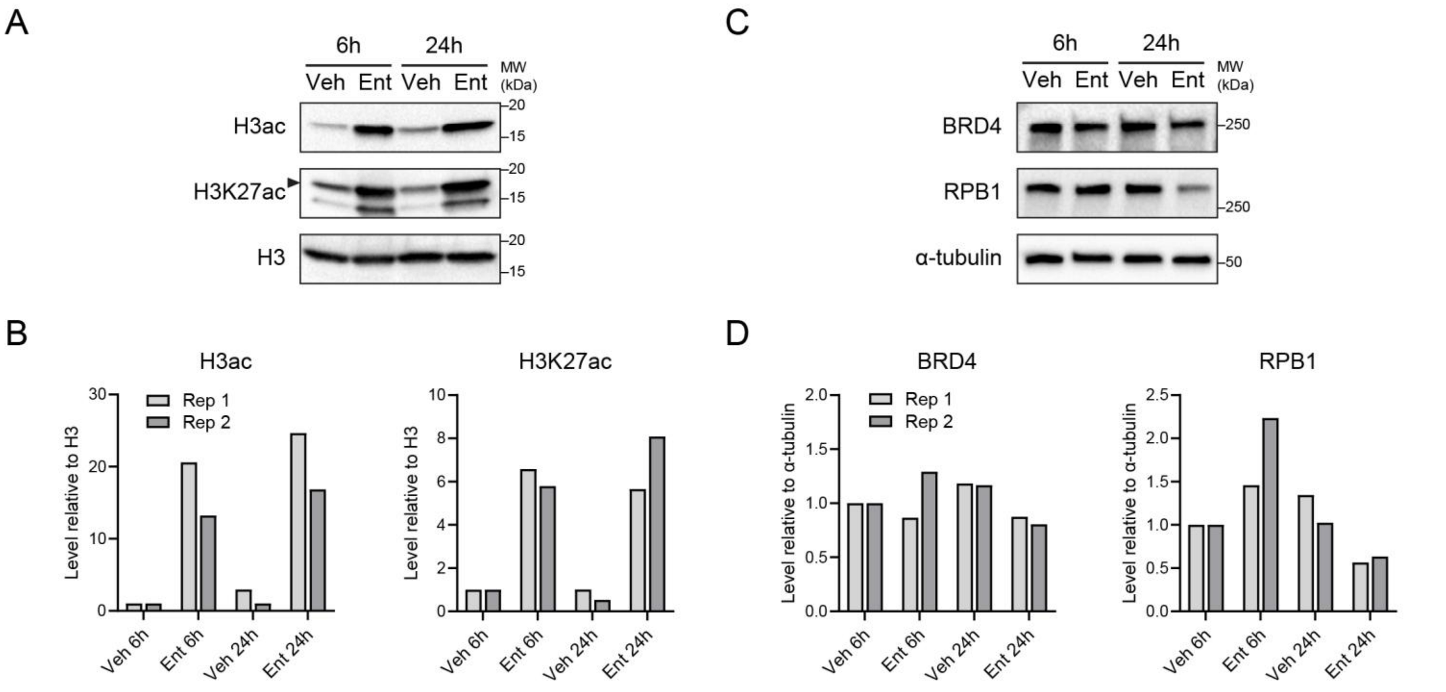


**Fig. S7.** Protein levels of H3 acetylation (H3ac), H3K27ac, BRD4, and RPB1 under entinostat treatments. (*A–D*) Representative Western blots and quantifications showing global H3ac and H3K27ac (*A*, *B*; normalized to H3) and BRD4 and RPB1 (*C*, *D*; normalized to α-tubulin) in FC1245 cells treated with Ent (5 μM, 6 or 24 h). *n* = 2 independent experiments.

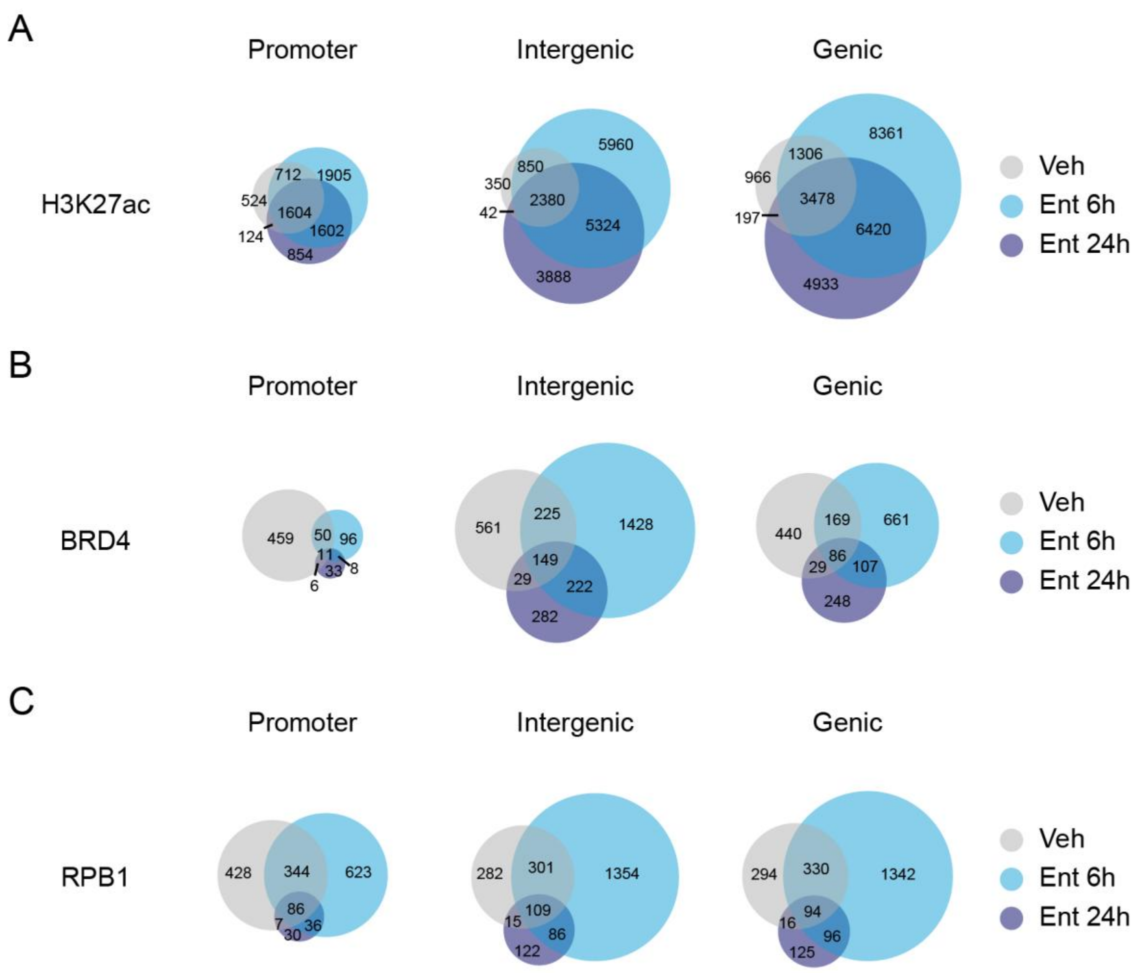


**Fig. S8.** Entinostat treatment redistributes H3K27ac, BRD4, and RPB1 peaks in individual annotation categories. (*A*–*C*) Venn diagrams of H3K27ac (*A*), BRD4 (*B*), and RPB1 (*C*) peak distributions across treatments by annotation categories.

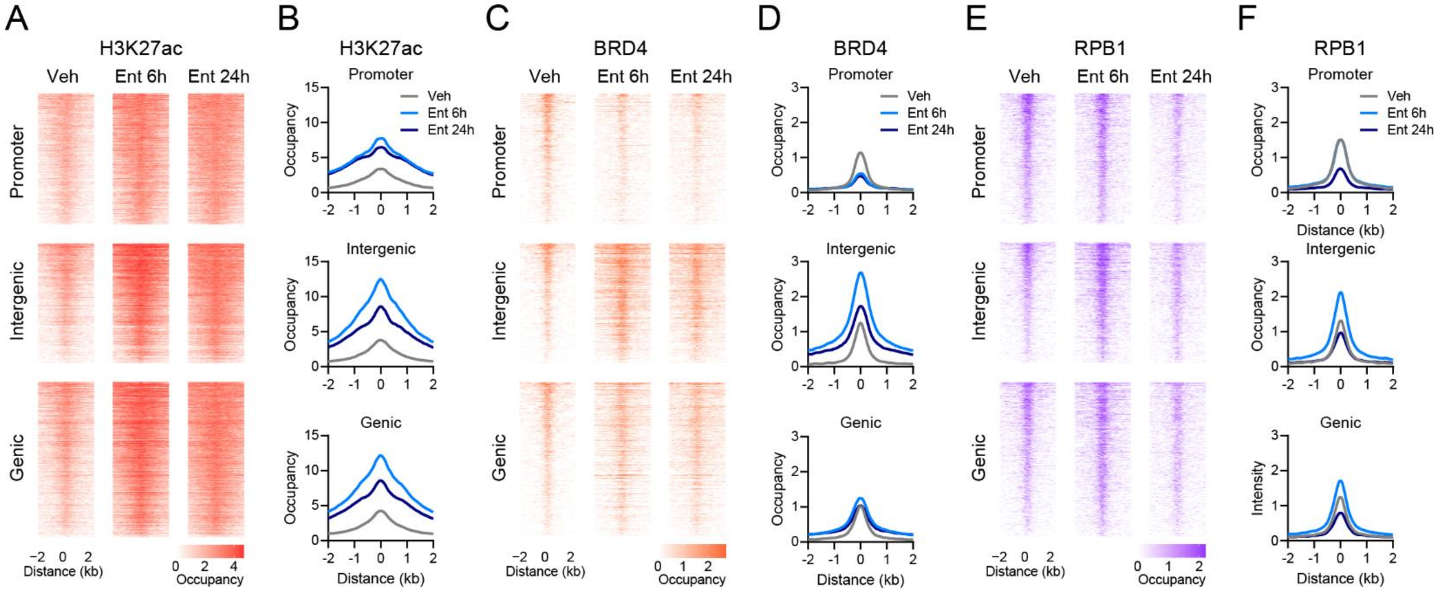


**Fig. S9.** Genomic location-dependent dynamics of H3K27ac, BRD4, and Pol II at HDAC1/2-bound sites. (*A*–*F*) Heatmaps and average profiles of H3K27ac (*A*, *B*), BRD4 (*C*, *D*), and RPB1 (*E*, *F*) occupancy at HDAC1/2-bound promoter, intergenic, and genic sites.

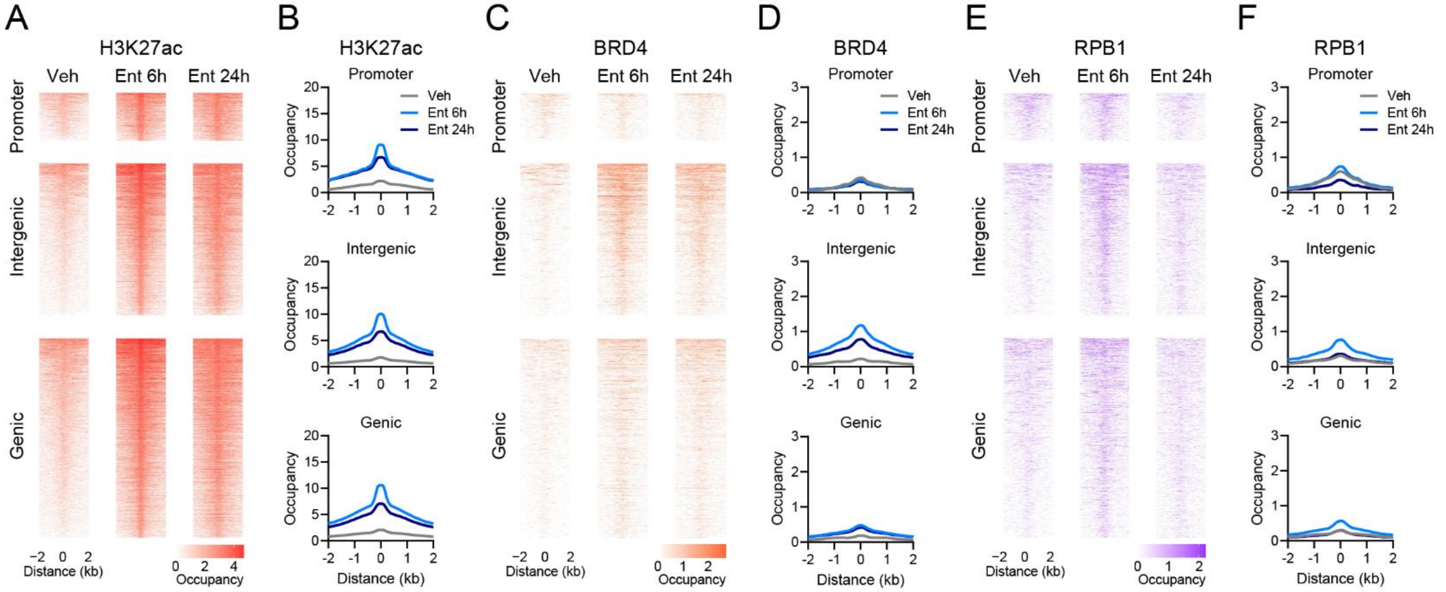


**Fig. S10.** Entinostat-induced intergenic acetylation gain recruits transcription machinery at de novo H3K27ac sites. (*A*–*F*) Heatmaps and average profiles of H3K27ac (*A*, *B*), BRD4 (*C*, *D*), and RPB1 (*E*, *F*) occupancy at promoter, intergenic, and genic H3K27ac sites gained after 6 h of Ent treatment.

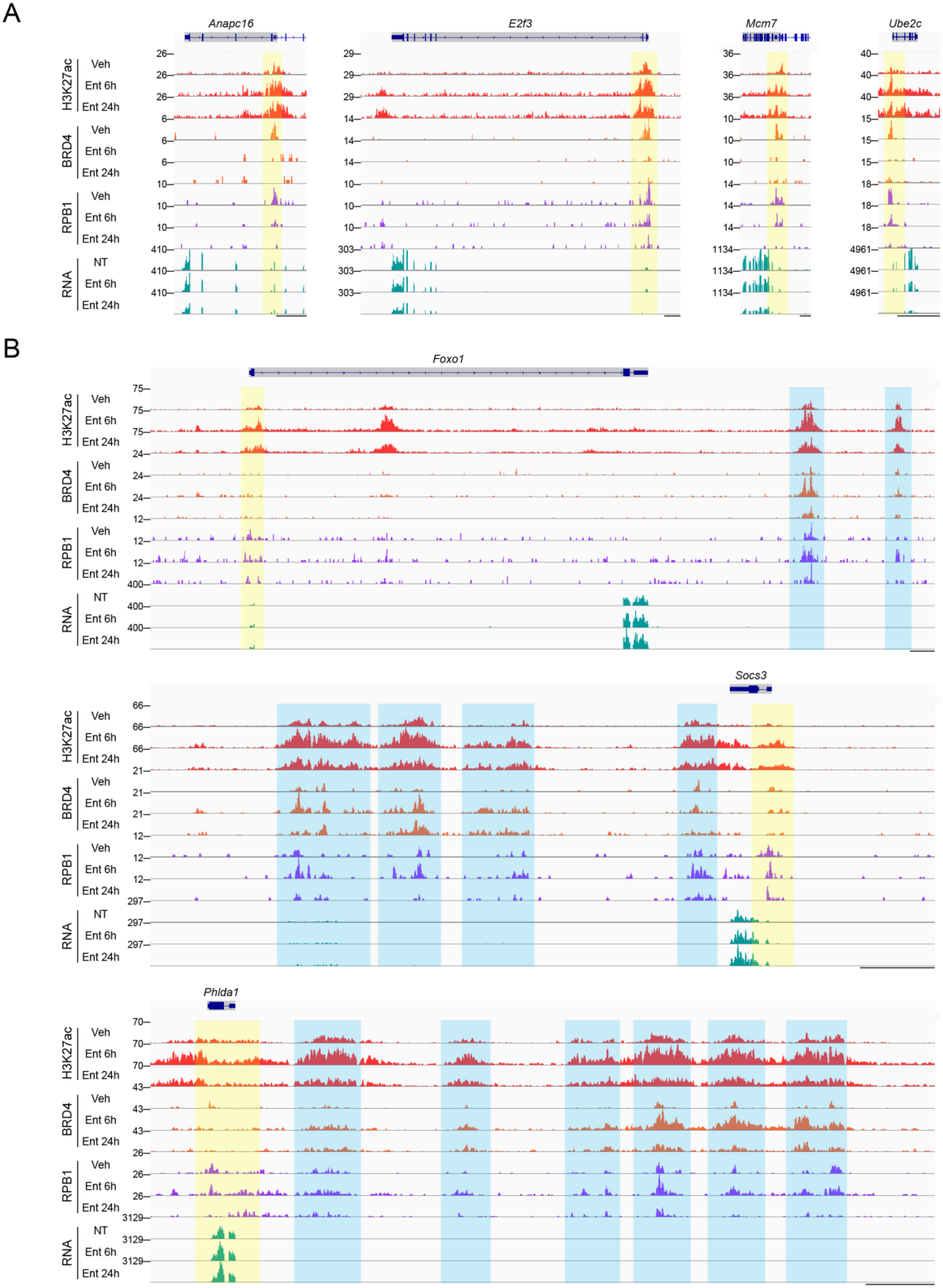

A
Anapc16
E2f3
Mcm7
Ube2c
H3K27ac
BRD4
RPB1
RNA
Veh
Ent 6h
Ent 24h
NT
26
6
10
410
29
14
303
36
1134
40
15
18
4961
B
Foxo1
75
24
12
400
Socs3
66
21
297
Phlda1
70
43
26
3129

**Fig. S11.** Transcription machinery dynamics at entinostat-regulated gene loci upon HDAC inhibition. (*A* and *B*) Genome browser tracks showing H3K27ac, BRD4, and RPB1 dynamics at representative entinostat-downregulated cell cycle genes (*A*) and entinostat-upregulated genes associated with differentiation (*Foxo1*, *Socs3*) and/or apoptosis (*Foxo1*, *Phlda1*) (*B*), together with corresponding RNA output. Promoters are highlighted in yellow and associated intergenic regions in blue. Scale bar, 5 kb.

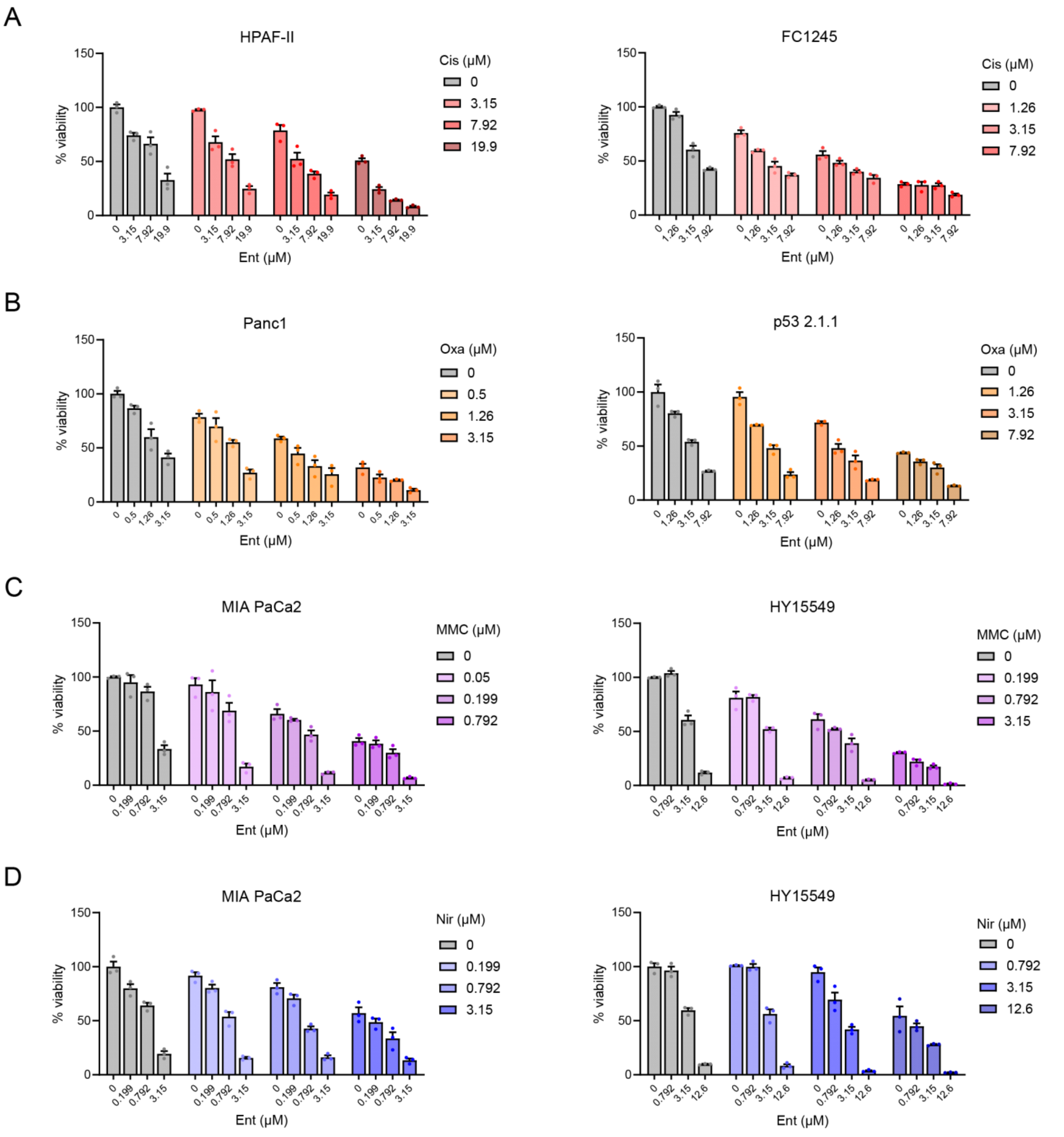


**Fig. S12.** Dosage-dependent reduction of PDAC cell viability under co-treatments of Ent with DNA damage-inducing agents. (*A–D*) Cell viability in representative human and mouse PDAC cell lines under graded combinations of Ent and Cis (*A*), Oxa (*B)*, MMC (*C*), or Nir (*D*). *n* = 3 cell sample replicates. Data are presented as mean values ± SEM.

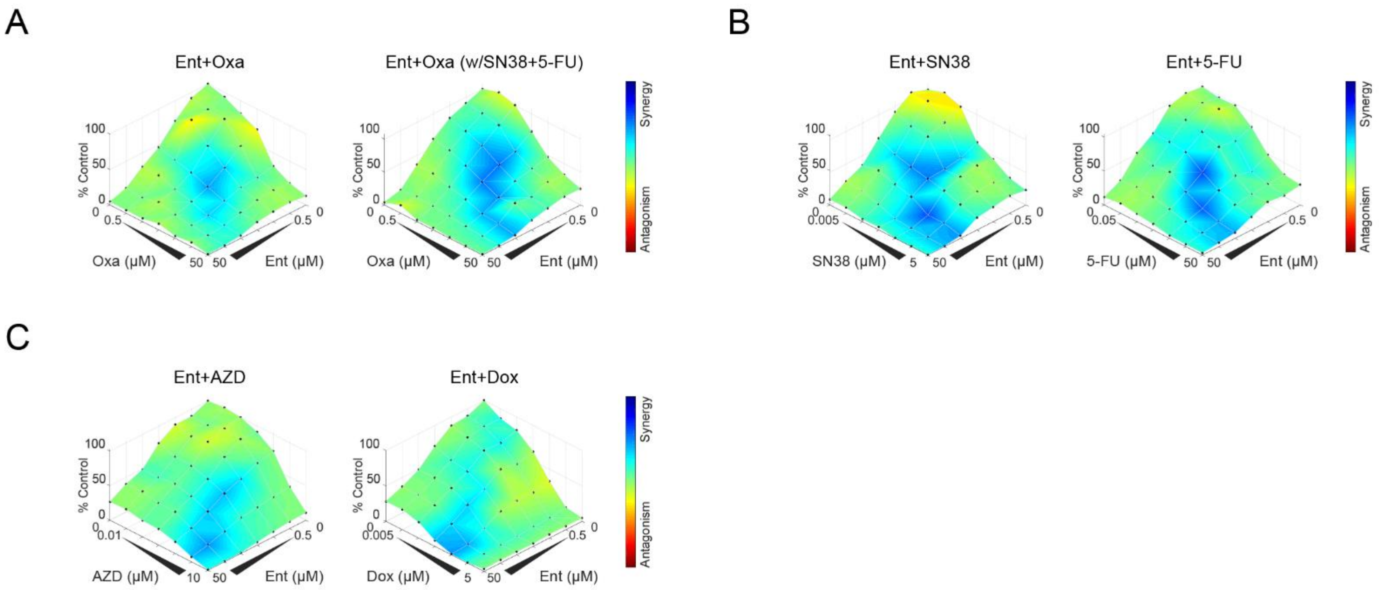


**Fig. S13.** Entinostat enhances the efficacy of diverse agents inducing DNA damage in vitro. (*A*) Synergy plots of viability in FC1245 cells under graded co-treatments of Ent and Oxa in the absence (*left*) or presence (*right*) of SN38 (0.02 µM) and 5-FU (0.8 µM). (*B* and *C*) Synergy plots of viability in FC1245 cells under graded co-treatments of Ent and SN38 (*B, left*), 5-FU (*B*, *right*), AZD0156 (AZD; *C*, *left*), or doxorubicin (Dox; *C*, *right*). *n* = 3 cell sample replicates.

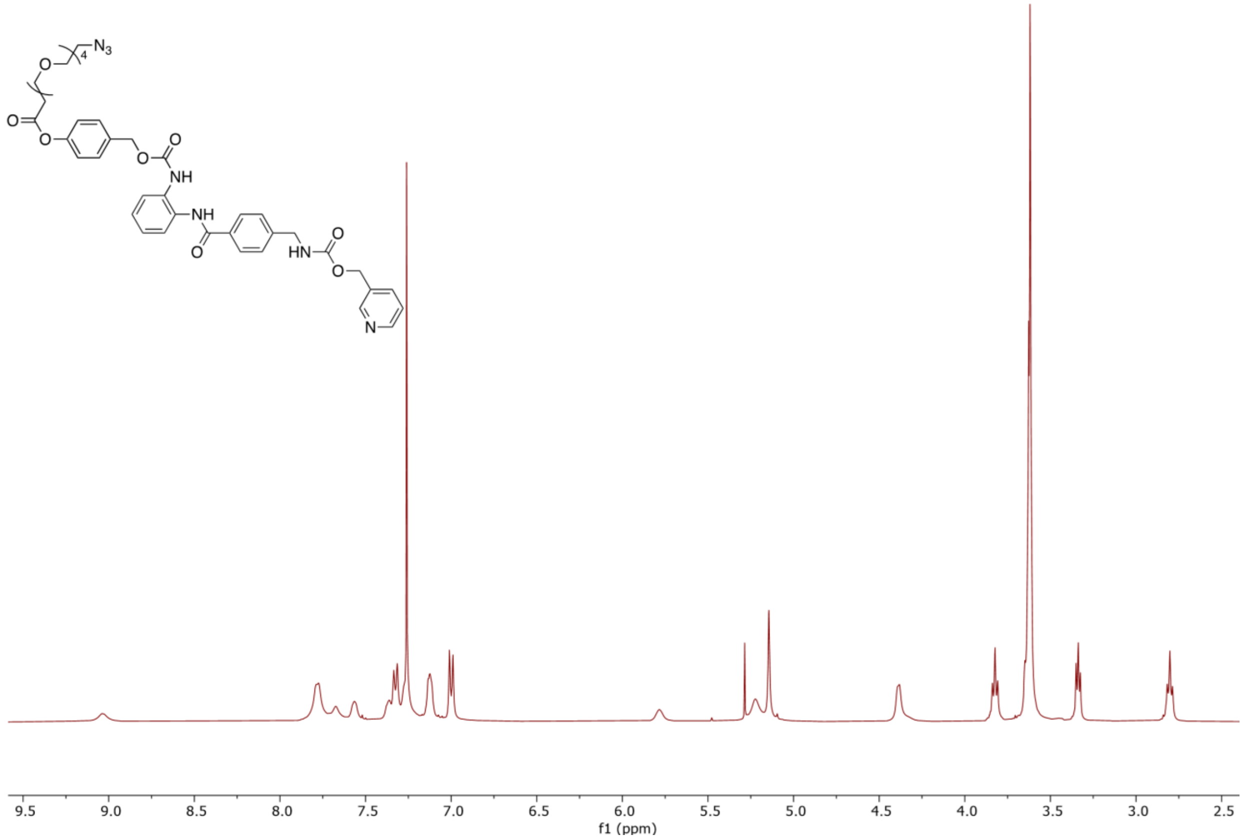


**Fig. S14.** $^{1}H$ NMR spectrum of Compound 2 in $CDCl_3$.

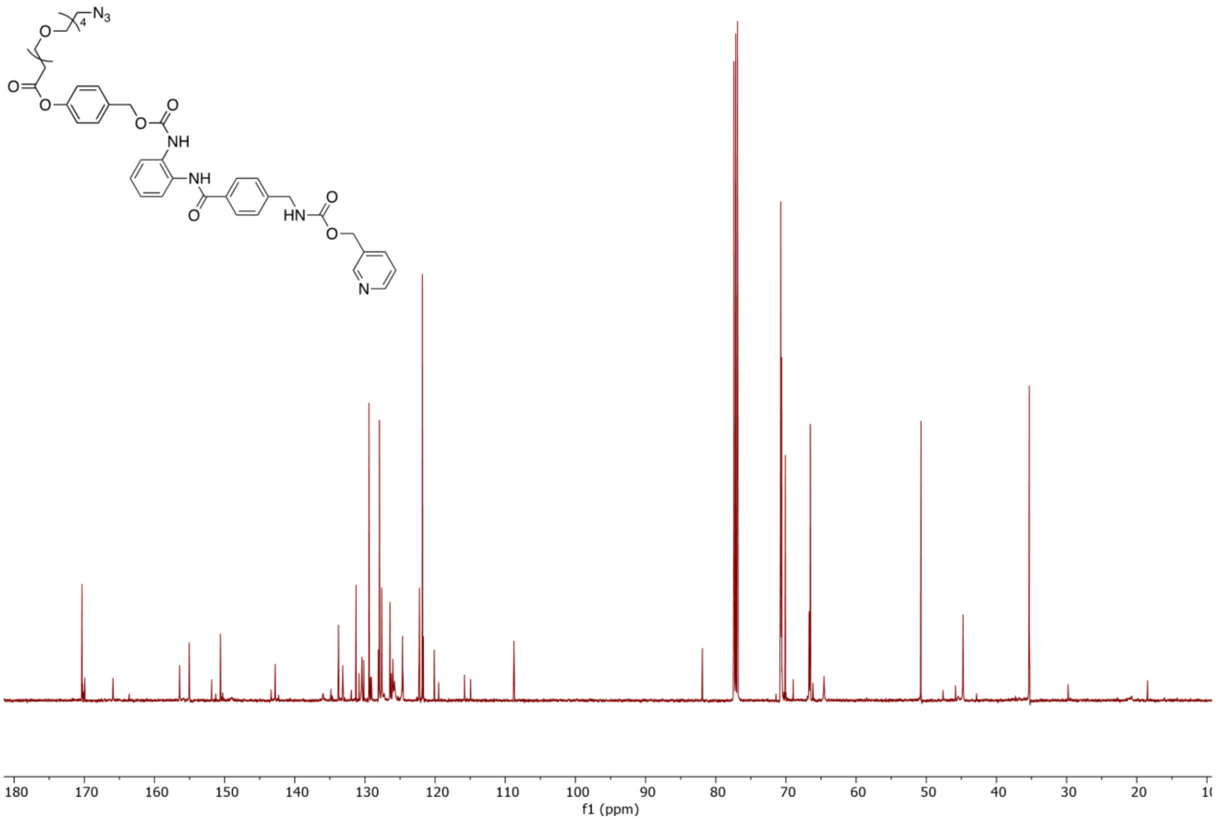


**Fig. S15.** $^{13}$C NMR spectrum of Compound 2 in $CDCl_3$.

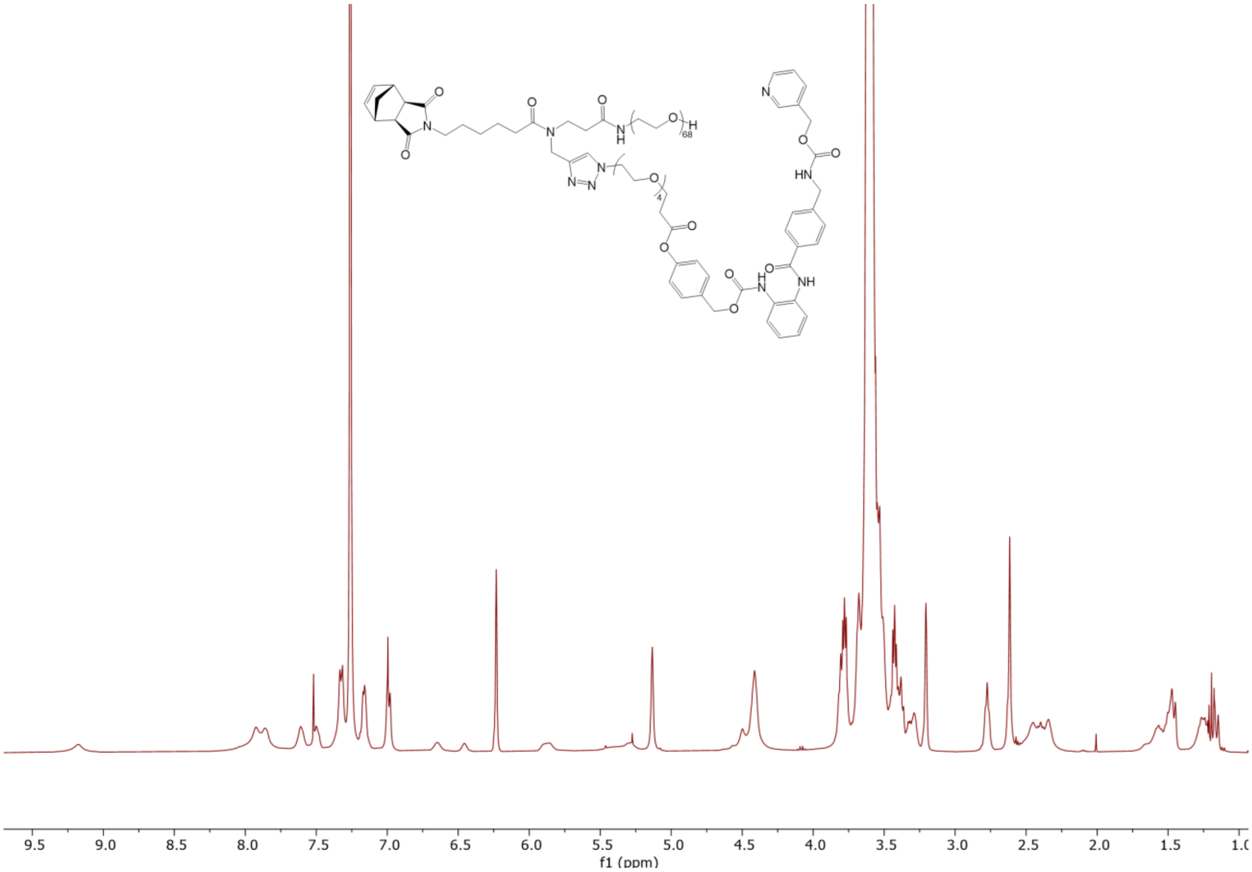


**Fig. S16.** $^{1}$H NMR spectrum of Ent-M in $CDCl_3$.

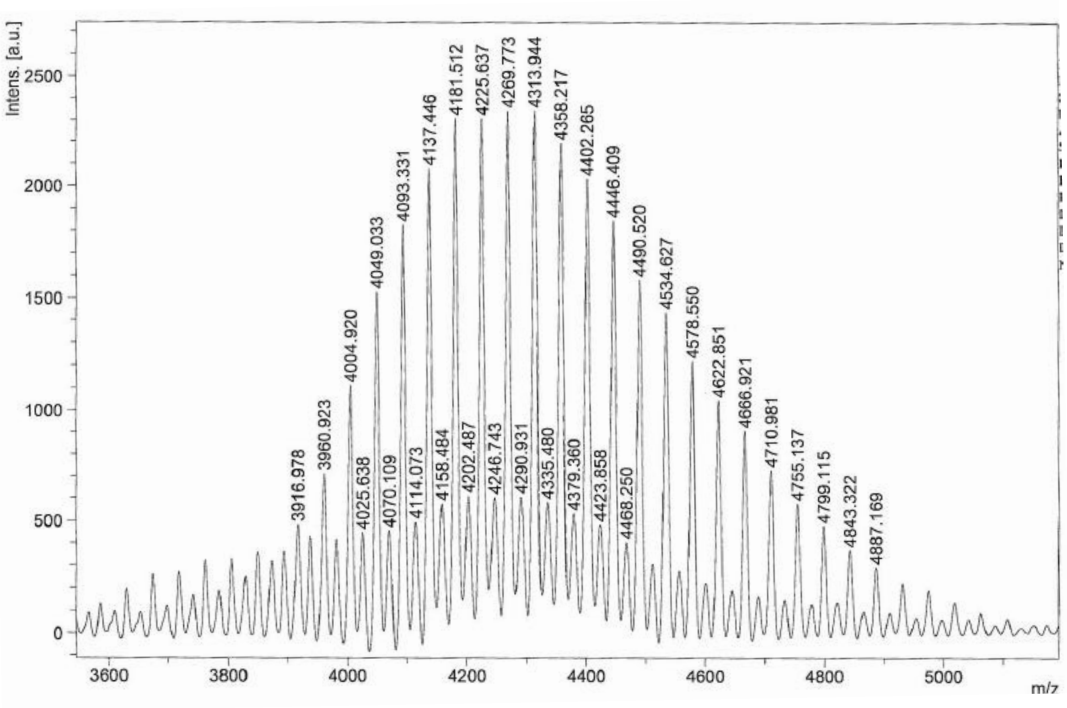


**Fig. S17.** MALDI-TOF spectrum of Ent-M. $C_{197}H_{345}O_{83}N_{10}$: m/z calculated 4181.91, found 4181.512 $[M + H]^+$. $C_{197}H_{344}O_{83}N_{10}Na$: m/z calculated 4203.30, found 4202.487 $[M + Na]^+$.

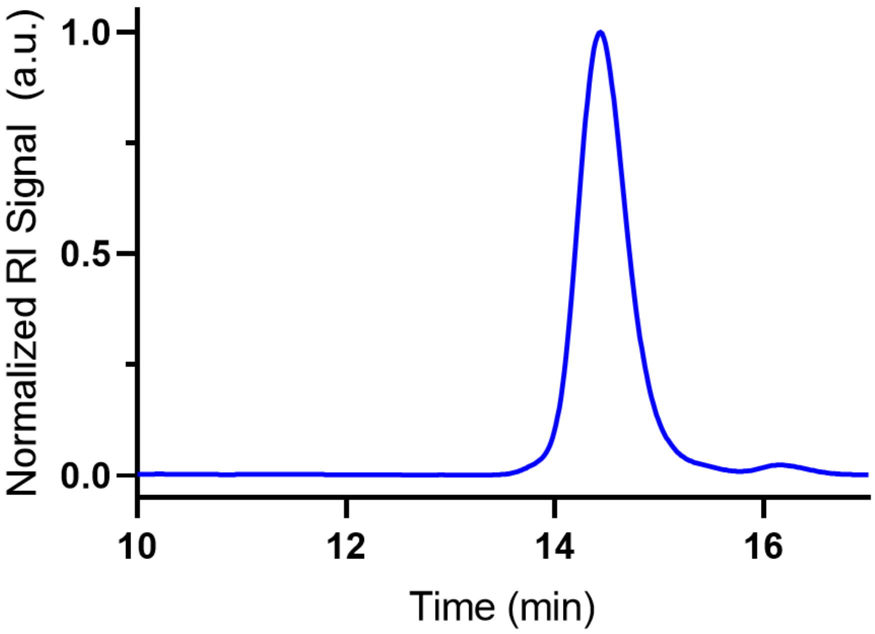


**Fig. S18.** GPC trace of Ent-BPD. Mw = 54.7 kDa; Mn = 49.6 kDa; Đ = 1.19. Molecular weight (MW) was acquired using a static light scattering detector. The minor peak at ~15 min elution time corresponds to residual Ent-M.

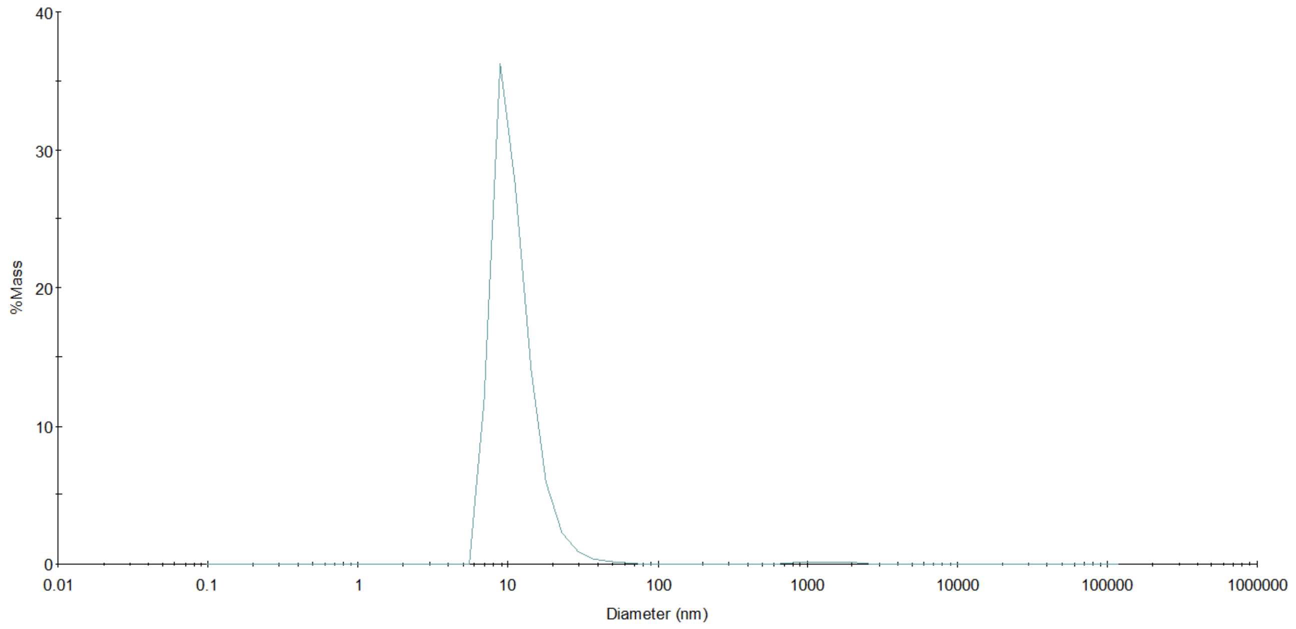


**Fig. S19.** Dynamic light scattering trace of Ent-BPD. Hydrodynamic diameter ($D_h$), as determined by DLS, to be 11 ± 5 nm.

**Table S1.** Diverse DNA damage-inducing agents evaluated for synergy with entinostat.

| Category | Class | Examined Drugs |
|---|---|---|
| DNA-damaging agents | Alkylating agents | Temozolomide |
| | Base analogs | 5-FU*, Gemcitabine |
| | Crosslinking agents, non-platinum-based | Mitomycin C* |
| | Crosslinking agents, platinum-based | Cisplatin*, Oxaliplatin* |
| | Intercalating agents | Doxorubicin* |
| | Radiomimetics | Bleomycin |
| | Topoisomerase I inhibitors | SN38* |
| | Topoisomerase II inhibitors | Etoposide |
| DDR inhibitors | ATM inhibitors | AZD0156* |
| | ATR inhibitors | Berzosertib |
| | CHK1 inhibitors | Rabusertib |
| | DNA-PK inhibitors | Nedisertib |
| | PARP inhibitors | Niraparib* |
| | WEE1 inhibitors | Adavosertib |

* Drugs shown synergy with entinostat in selected PDAC cell line(s).

### MATERIALS AND METHODS

#### Cell Culture

Mouse PDAC cell line FC1245 (*Kras*$^{LSL-G12D/+}$; *Trp53*$^{LSL-R172H/+}$; *Pdx1-Cre,* C57BL/6J) (1, 2), HY15549 (*Kras*$^{LSL-G12D/+}$; *Trp53*$^{f/+}$; *Ptf1a-Cre*, C57BL/6) (3), and p53 2.1.1 (*Kras*$^{LSL-G12D/+}$; *Trp53*$^{f/+}$; *Ptf1a-Cre*, FVB/NJ) (3, 4) were provided by David Tuveson, Haoqiang Ying, and Eric Collison, respectively. Human PDAC cell lines HPAC, HPAF-II, MIA PaCa2, PSN1, and Panc1 were acquired from ATCC. Cells were cultured in DMEM (Corning) with 10% characterized fetus bovine serum (HyClone) and 1× Antibiotic/Antimycotic (Gibco).

#### In Vitro Drug Treatment

Stock solutions of entinostat (WuXi AppTec), SN38 (Selleck Chemicals, S4908), 5-FU (Torcis, 3257), mitomycin C (Selleck Chemicals, S8146), niraparib (LC Laboratories, N-5665), and doxorubicin (Torcis, 2252) were prepared in dimethylsulfoxide (DMSO) at 10 mM and stored at −20ºC. Cisplatin (Torcis, 2251) and oxaliplatin (Torcis, 2623) were dissolved in water containing 0.3% Tween-20 at 5 mM and stored at room temperature and 4 ºC, respectively, for up to a month. AZD0156 (Selleck Chemicals, S1525) was prepared in DMSO/ethanol (1:1, v/v) at 2 mM and stored at −20ºC. Working concentrations were prepared immediately before treatment, manually or using a digital dispenser (HP, D300e).

#### shRNA-Mediated Depletion

An shRNA co-targeting *Hdac1* and *Hdac2* was expressed in the lentiviral vector pTY-U6-Pgk-Puro, with the seed sequence of 5'-GCAAGCAGATGCAGAGATTCA-3'. Individual shRNAs targeting *Hdac1*, *Hdac2*, or *Hdac3*, as well as the procedure of lentivirus production, were described previously (5). Transduced cells were subjected to puromycin selection (2 µg/ml) for 4 d prior to RNA isolation.

#### In Vitro Viability Assay

PDAC cells were seeded in 384-well plates with 30 µl of culture medium each well, 24 h before drug treatment. Seeding density and treatment duration were optimized for each cell line according to proliferation rate: FC1245, 150 cells/well, 3 d; HY15549, 120 cells/well, 3 d; p53 2.1.1, 120 cells/well, 2 days; HPAC, 420 cells/well, 4 d; HPAF-II, 600 cells/well, 4 d; MIA PaCa2, 300 cells/well, 3 d; Panc1, 420 cells/well, 4 d; PSN1, 600 cells/well, 3 d. Cell viability was measured at treatment endpoints using CellTiter-Glo assay (Promega, G9242). Viability was calculated relative to untreated controls, and drug synergy analyzed with Combenefits using the highest single agent (HSA) model (6).

### RNA-seq Analysis

Total RNA was extracted with the RNeasy Mini Kit, assessed for quality with a Bioanalyzer (Agilent, 2100), and used to generate cDNA libraries with TruSeq Stranded RNA Sample Preparation Kit (Illumina, v2). Libraries were pooled and sequenced on the HiSeq 2500 system (Illumina) with single-end 100-bp reads. Reads were aligned to UCSC reference genomes mouse (mm10 or 39) or human reference genomes (hg19) and quantified with Salmon (7). Differential expression analysis was performed with DESeq2 (8), and temporal dynamics with Mfuzz (v2.64.0) (9). GSEA and GO analysis was performed using GSEA (Broad Institute, v4.0.3) (10) and DAVID (v6.8) (11), respectively.

### CUT&RUN Analysis

Tumor cell samples ($10^5$ cells each) were subjected to CUT&RUN (Cell Signaling, 86652) with antibodies against HDAC1 (Cell Signaling, 34589), HDAC2 (Abcam, ab32117), H3K27ac (Cell Signaling, 8713), BRD4 (EpiCypher, 13-2003), and RPB1 (Cell Signaling, 2629). Genomic DNA from *Saccharomyces cerevisiae* was added as a spike-in control for sample normalization. Purified DNA fragments were used to construct libraries with the TruSeq ChIP Library Preparation Kit (Illumina, IP-202-1012). Libraries were sequenced as paired-end 100-bp reads on a NovaSeq 6000 System (Illumina). Reads were aligned to UCSD mouse (mm39) and yeast (sacCer3) reference genomes using STAR (v2.7.11b) (12). Mouse-specific reads were normalized to yeast-specific read count in each sample. PCR duplicates, reads mapping to both genomes, mitochondrial reads, and reads with ambiguous chromatin locations were removed. Peak calling was performed with MACS3 (v3.0.1) (13), and downstream peak and motif analyses with HOMER (UCSD, v4.0) (14). Annotated peaks were classified into genic (3'-UTR, 5'-UTR, exon, intron, TTS), intergenic, promoter (/TSS), and other (miRNA, ncRNA, rRNA, scRNA, snoRNA, snRNA, pseudogene).

### Western Blotting

Protein extracts were prepared in RIPA buffer from cells lysed on plates or from tissues homogenized by a PowerLyzer (Qiagen). Samples were subjected to SDS-PAGE and Western blotting with primary antibodies against γH2A.X (Cell Signaling, 2577; 1:1000), H2A.X (Cell Signaling, 2595; 1:1000), H3ac (histone H3 acetyl-K9/K14/K18/K23/K27; Abcam, ab47915; 1:5000), H3K27ac (Abcam, ab4729; 1:1000), H3 (Abcam, ab1791; 1:1000), BRD4 (Bethyl, A301-985A50; 1:2000), RPB1 (Active Motif, 39097; 1:5000) and α-tubulin (Sigma, T6199; 1:1000), as well as HRP-conjugated secondary antibodies against rabbit (Cell Signaling, 7074; 1:5000) and mouse IgG (Santa Cruz, sc-2005; 1:5000) were used. Images were acquired with a ChemiDoc XRS+ system (Bio-rad) and quantified using Image Lab software (Bio-Rad, v5.2.1).

### Animal Housing

Mice were housed at a temperature (22±1 °C) and humidity (45–65%) controlled environment under a 12 h light-dark cycle with ad libitum access to food and water.

### Orthotopic Transplantation

Wild-type C57BL/6J male mice (8-12 weeks; the Jackson Laboratory) were used for orthotopic transplantation. Mouse PDAC cells, FC1245 (100 cells) or HY15549 (200 cells), suspended in Matrigel (1:1, v/v), were injected into the pancreatic parenchyma. Mice were enrolled for treatment either 10 d post-transplantation or upon detection of sizable tumors (~60 $mm^3$) by ultrasound imaging. Tumor burden was assessed by endpoint tumor weight or ultrasound-based tumor volume ($0.52 \times x \cdot \times y \times z$). No tumors exceeded 2,000 $mm^3$, the IACUC-approved limit.

### In Vivo Administration of Small-Molecule Drugs and BPDs

Entinostat was dissolved in phosphate-buffered saline (PBS) with 0.05 N HCl and 0.1% Tween-20 at 5 mg/ml, sterile-filtered, and administrated by oral gavage. Cisplatin and oxaliplatin were dissolved in saline at 0.5 mg/ml, sterile-filtered, and administrated by intraperitoneal injection. Entinostat-BPD and Cy7.5-BPD were dissolved in PBS and administrated by intraperitoneal injection.

### Whole Blood Collection and Analysis

Whole blood was collected by cardiac puncture at endpoint under terminal anesthesia. Hematological analysis was performed by IDEXX BioAnalytics.

### Ex Vivo Fluorescence Imaging

Tumors, tissues, and organs harvested from orthotopic models 24 h after Cy7.5-BPD administration were imaged using an IVIS Spectrum Imaging System and quantified with Living Image software (Caliper Life Sciences).

### IHC Analysis

Tumors were fixed in 10% buffered formalin overnight at 4 °C, transferred to 70%, and paraffin embedded. Sections (5 µm) were subjected to antigen retrieval in citric acid-based buffer (Vector Labs, H-3300) using a pressure cooker for 15 min. IHC was performed with anti-γH2A.X antibody (Cell Signaling, 9718; 1:480) and the staining kit (Cell Signaling, 13079).

Slides were scanned on an Axioscan 7 (Zeiss) or VS-100 (Olympus) scanning microscope and quantified with QuPath (v0.4.0).

### Synthesis of Entinostat-BPD (Ent-BPD)

#### *Section A. Materials, General Methods, & Instrumentation*

**Materials.** All reagents were purchased from commercial suppliers and used without further purification unless otherwise stated. Grubbs 3rd generation bispyridyl catalyst **G3-cat** (15), linker precursor **Compound 1** (16), and macromonomer precursor **yne-M** (17) were prepared as previously reported. Column chromatography was carried out on silica gel 60F (EMD Millipore, 0.040–0.063 mm) or on aluminum oxide (Sigma-Aldrich, activated, neutral, Brockmann Activity I). Recycling preparative HPLC was performed on a LaboACE system (Japan Analytical Industry), using a JAIGEL-2 HR JAIGEL-2.5HR column in series.

**Gel permeation chromatography (GPC)**. Analyses were performed on an Agilent 1260 Infinity setup with two Agilent PL1110-6500 columns in tandem and a 0.025 M LiBr DMF mobile phase run at 60°C. Data collection was done using Wyatt ASTRA 6.1 and plotted using GraphPad Prism 8.0.2. The differential refractive index (dRI) of each compound was monitored using a Wyatt Optilab T-rEX detector; the light scattering (LS) signal was acquired with a Wyatt Dawn Heleos-II detector.

**Nuclear magnetic resonance (NMR).** Spectra were recorded on a Bruker AVANCE III-400 spectrometer with working frequencies of 400 ($^{1}$H) and 100 ($^{13}$C) MHz or on an AVANCE-500 spectrometer with working frequencies of 500 ($^{1}$H) and 126 ($^{13}$C) MHz. Data collection was done using Bruker Topspin 4.0, and analyses were done with MestReNova (v12.0.4). Chemical shifts are reported in ppm relative to the signals corresponding to the residual non-deuterated solvents: $CDCl_3$: δH = 7.26 ppm and δC = 77.16 ppm. High-resolution mass spectra (HRMS) were measured on a JEOL AccuTOF LC-Plus 4G with an IonSense DART and analyzed with msAxel (v1.0.5.2). Matrix-assisted laser desorption/ionization time-of-flight (MALDI-TOF) analyses were collected with Bruker Daltonics FlexAnalysis 3.4 on a Bruker OmniFlex instrument, using sinapinic acid as the matrix.

**Dynamic light scattering (DLS).** Measurements were performed on a Wyatt Technology Mobius DLS instrument. Data collection was done using Wyatt Dynamics 7.5.0.17. Nanoparticle suspensions were prepared in a solution of nanopure water (MilliQ), PBS buffer, or 5% w/w glucose/nanopure water (1 mg/ml). The resulting suspensions were passed through a 0.45 μm Nalgene filter (PES membrane) into disposable polystyrene cuvettes, which were pre-cleaned with compressed air. Measurements were made in sets of 10 acquisitions, and the average hydrodynamic diameters were calculated by using the DLS correlation function via a regularization fitting method (Dynamics 7.5.0.17 software).

### *Section B. Synthetic Procedure of Ent-BPD*

#### 1. Small Molecule Precursor

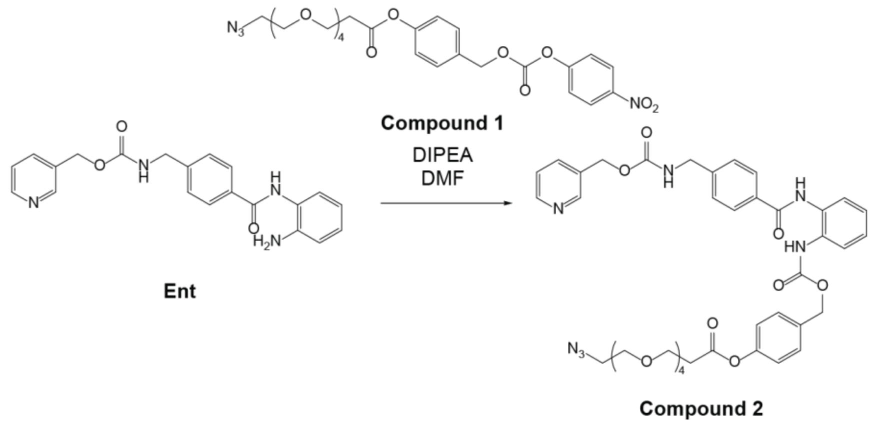


**Synthesis of Compound 2.** To an 8 ml vial, **Ent** (111 mg, 295 μmol, 1.07 eq), **Compound 1** (133 mg, 272 mmol, 1.0 eq), DIPEA (21.4 mg, 28.8 μL, 166 μmol, 0.61 eq), HOBt (33.6 mg, 249 μmol, 0.92 eq), pyridine (238 μL) and DMF (8.0 ml) were added. The reaction mixture was allowed to stir for 48 h at rt. EtOAc was then added, and the reaction mixture was washed with water (X2) and brine (X1). The organic layer was collected, dried over $Na_2SO_4$, filtered, and concentrated under vacuum. The resulting mixture was redissolved in $CHCl_3$, filtered through a 0.45 μm filter, and subjected to recycling preparative HPLC. The fractions containing the product were concentrated under vaccum and dried overnight, affording the pure product as a solid (107 mg, 57% yield). See Figs. S9 and S10 for the $^1H$ and $^{13}C$ NMR spectra of **Compound 2**. LRMS: Calcd for $C_{40}H_{46}N_7O_{11}$: m/z = 800.33 $[M + H]^+$; Found: 800.1. $^1H$ NMR (400 MHz, CDCl3) δ 9.04 (s, 1H), 7.89 – 7.49 (m, 5H), 7.39 – 7.21 (m, 7H), 7.18 – 7.06 (m, 2H), 7.00 (d, J = 8.0 Hz, 2H), 5.78 (s, 1H), 5.31 –5.08 (m, 5H), 4.39 (s, 2H), 3.82 (t, J = 6.2 Hz, 2H), 3.74 – 3.48 (m, 14H), 3.34 (t, J = 5.1 Hz, 2H), 2.80 (t, J = 6.2 Hz, 2H). $^{13}C$ NMR (126 MHz, $CDCl_3$) δ 170.33, 156.42, 155.04, 150.60, 142.79, 133.78, 133.16, 131.28, 130.44, 130.18, 129.42, 128.10, 127.93, 127.62, 126.43, 124.65, 122.25, 121.81, 121.79, 121.67, 120.13, 108.77, 81.91, 70.79, 70.77, 70.76, 70.74, 70.70, 70.61, 66.70, 66.52, 66.49, 50.78, 50.76, 44.76, 35.33.

#### 2. Macromonomer

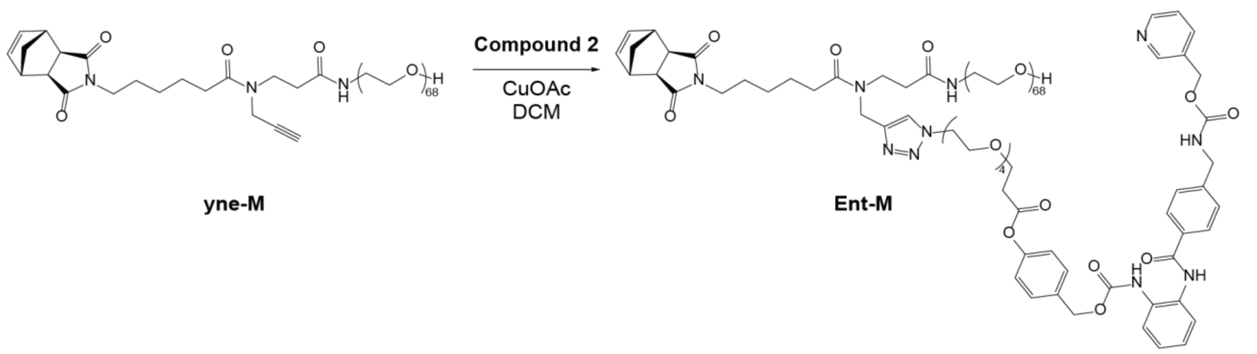


**Synthesis of Ent-Loaded Macromonomer (Ent-M).** To a vial, **yne-M** (152 mg, 45.1 µmol), **2** (47.0 mg, 57.7 µmol), and DCM (10 ml) were added. CuOAc (a pinch) was then added, and the reaction mixture was stirred under N2 atmosphere. The reaction was complete in ~1 h as determined by LC-MS. The crude mixture was run through an aluminum oxide plug. The collected solution was concentrated under vacuum, redissolved in $CHCl_3$, filtered through a 0.45 µm filter, and subjected to recycling preparative HPLC. The fractions containing the product were concentrated under vacuum and dried overnight, affording the product as a solid (161 mg, 85% yield). See Figs. S11 and S12 for $^1$H NMR and MALDI-TOF spectra of **Ent-M**.

### 3. Macromolecular BPD

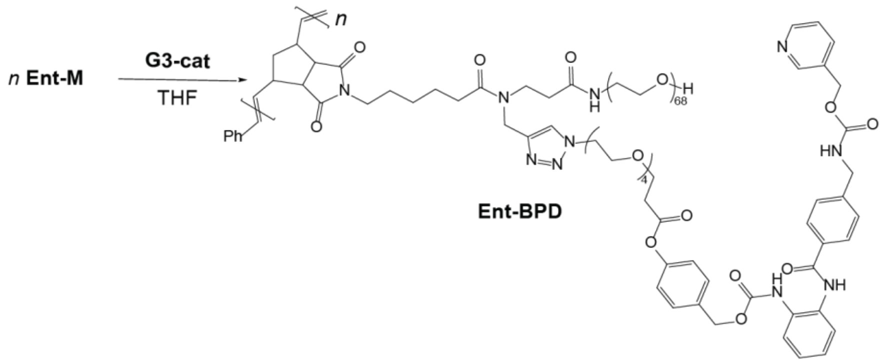


**Synthesis of Ent-BPD**. To a vial containing a stir bar, **Ent-M** (19.3 mg, 4.63 µmol, 10.0 eq) was added. To another vial, a solution of 3rd generation Grubbs catalyst (**G3-Cat**, 0.02 M in THF) was freshly prepared. THF (92.6 µl) was then added to the vial containing **Ent-M** followed by the addition of the **G3-Cat** solution (23.2 µl, 0.463 µmol, 1.0 eq) in order to give the desired degree of polymerization (DP) of 10 and to achieve a total macromonomer concentration of 0.05 M. The reaction mixture was allowed to stir for 6 h at room temperature. To quench the polymerization, a drop of ethyl vinyl ether was then added. The reaction mixture was transferred to 8 kDa molecular weight cutoff dialysis tubing in 3 ml nanopure water, and the solution was dialyzed against $H_2O$ (500 ml × 3, solvent exchange every 6 h). The dialyzed solution of **Ent-BPD** was then subjected to lyophilization. Alternatively, **Ent-BPD** was also acquired by precipitation in diethyl ether. See Figs. S13 and S14 for GPC and DLS characterization of **Ent-BPD**.
